# Lipids are essential for potassium transport by KdpFABC from E. coli

**DOI:** 10.64898/2026.03.20.713019

**Authors:** Adel Hussein, Xihui Zhang, Michael Schlame, Bjørn P. Pedersen, David L. Stokes

## Abstract

KdpFABC is a hetero-tetrameric potassium pump that uses ATP to import potassium and thereby maintain homeostasis in bacteria under stress conditions. KdpA is a channel-like subunit with a selectivity filter that binds potassium from the periplasm. K^+^ then moves through a ∼40Å-long intramembrane tunnel to reach a canonical binding site in KdpB. KdpB is a P-type ATPase that orchestrates conformational changes associated with the Post-Albers reaction cycle, involving E1 and E2 conformations and formation of an aspartyl phosphate intermediate as a way of coupling ATP hydrolysis to K^+^ transport. To elucidate the associated structural changes in a lipid environment, we reconstituted wild-type KdpFABC into lipid nanodiscs and used cryo-EM to image the complex under active turnover. The resulting six high resolution (2.1-2.7 Å) structures provide new insight into the sequence of allosteric changes that produce (1) occlusion of K^+^ at the canonical binding site and (2) expulsion of K^+^ from this site and into a low-affinity release site. The structures also reveal two types of lipids bound to the complex. Specifically, two structural lipids bind at subunit interfaces and ∼20 annular lipids are seen at the periphery of the complex. In addition, we tested functional effects of mutations to residues at the KdpA/KdpB interface. ATPase and transport assays were used to document functional defects that reflect delipidation of structurally compromised complexes. We conclude that lipids play an integral role in structure and function of the KdpFABC complex.

**Significance:** KdpFABC uses ATP to transport potassium across the plasma membrane of E. coli. To further our understanding of its mechanism, we put purified KdpFABC molecules into membrane bilayers and used cryo-EM to capture structures during active transport. We have thereby produced structures representing all major states of the transport cycle with a high degree of precision. Analysis of these structures reveals new details about two key steps in this cycle and shows lipid molecules bound to the protein. We then introduced mutations at the interface between the two main subunits, which controls passage of potassium across the membrane. Activity measurements reveal how the protein depends on lipid to stabilize the structure and facilitate transport.

## Introduction

KdpFABC is an ATP dependent membrane transporter that maintains potassium homeostasis under stress (1). It has been the subject of numerous studies since the 1970’s (2, 3), with mechanistic insight resulting from recent structural studies (4–9). The proposed transport mechanism starts with periplasmic K^+^ binding with high affinity to a classical ion selectivity filter in the channel-like KdpA subunit (10), which is a descendant from the Superfamily of K^+^ Transporters (11). Ions are then shuttled into a long, water-filled intramembrane tunnel leading to a canonical ion binding site (CBS) in the pump-like subunit, KdpB, which is a descendant of P-type ATPases (12). Arrival of K^+^ at the CBS triggers ATP hydrolysis by the cytoplasmic domains of KdpB, thus transitioning from the resting E1 state to the high-energy E1∼P state in which a conserved aspartate (Asp307) is auto-phosphorylated. The energy of this phosphoenzyme drives a major conformational change to the so-called E2-P state, which closes the tunnel and moves the K^+^ into a low-affinity exit site, from which it is released to the cytoplasm (7, 8). This mechanism conforms to the classical Post-Albers scheme employed by other P-type ATPases (13, 14) and provides the current framework for understanding transport by KdpFABC. Central questions remain about the energetics of moving K^+^ ions through the narrow, dewetted portion of the tunnel at the interface between KdpA and KdpB and allosteric mechanisms that couple ATP hydrolysis with ion movement (15). In addition, lipids have been shown to affect activity (5, 16) and a lipid molecule has been observed at a consistent site in previous structures from detergent-solubilized complexes (4–6, 8), suggesting that the membrane environment may be important for function.

For the current work, we reconstituted purified KdpFABC complexes into lipid nanodiscs and used cryo-EM to image samples under active turnover conditions. We previously reported on the 2.1 Å resolution structure of the most populated state corresponding to the E1∼P·ADP state (9). We have now resolved five additional conformational states with a variety of ATP-related ligands at the phosphorylation site with resolutions between 2.3 and 2.8 Å. Based on the nature of these ligands and on comparisons with previous structural results, we have associated each structure with a reaction intermediate from the Post-Albers catalytic cycle. These new structures confirm conformational dynamics previously seen in detergent solution, with higher resolutions that reveal new details associated with occlusion of the intramembrane tunnel and ion displacement from the CBS. Two lipid molecules are bound at subunit interfaces, one of which has been previously reported whereas the second appears to be novel. In addition, all six structures reveal non-protein densities around the periphery of the complex which we have modeled as ∼20 annular lipid molecules. We went on to explore the functional effects of mutations at the KdpA/KdpB subunit interface and at the pinch point of the intersubunit tunnel, thus revealing a variety of lipid-dependent effects on complex stability and on allosteric coupling between ATP hydrolysis and ion transport.

## Results

### Structures of KdpFABC under turnover conditions

As described in our recent report of the E1∼P·ADP structure (9), we reconstituted purified, detergent solubilized wild-type protein into lipid nanodiscs to elucidate structural details in a more native, membrane-bound state. Similar to an earlier study by Silberberg et al. (6), Mg·ATP was added to stimulate active turnover prior to cryo-EM sample preparation, thus generating a heterogenous population of particles representing all states from the Post-Albers transport cycle. Initial 3D classification of ∼50,000 images produced two main classes distinguished by the orientation of cytoplasmic domains (designated A, N and P) and consistent with E1 and E2 states (Sfig. 1). To segregate sub-states, we used focused 3D classification in which a mask was applied to the cytoplasmic domains of KdpB. This classification generated four E1 sub-states and two E2 sub-states (Fig. 1, Sfig. 1). Given their dynamic nature, the resolution of the cytoplasmic domains of KdpB was much lower. We therefore implemented masked refinement to improve the resolution in this region of the map (Sfig. 2). Final atomic models were produced using the globally refined map for the membrane domains and the map from masked refinement for the cytoplasmic domains.

**Figure 1.**
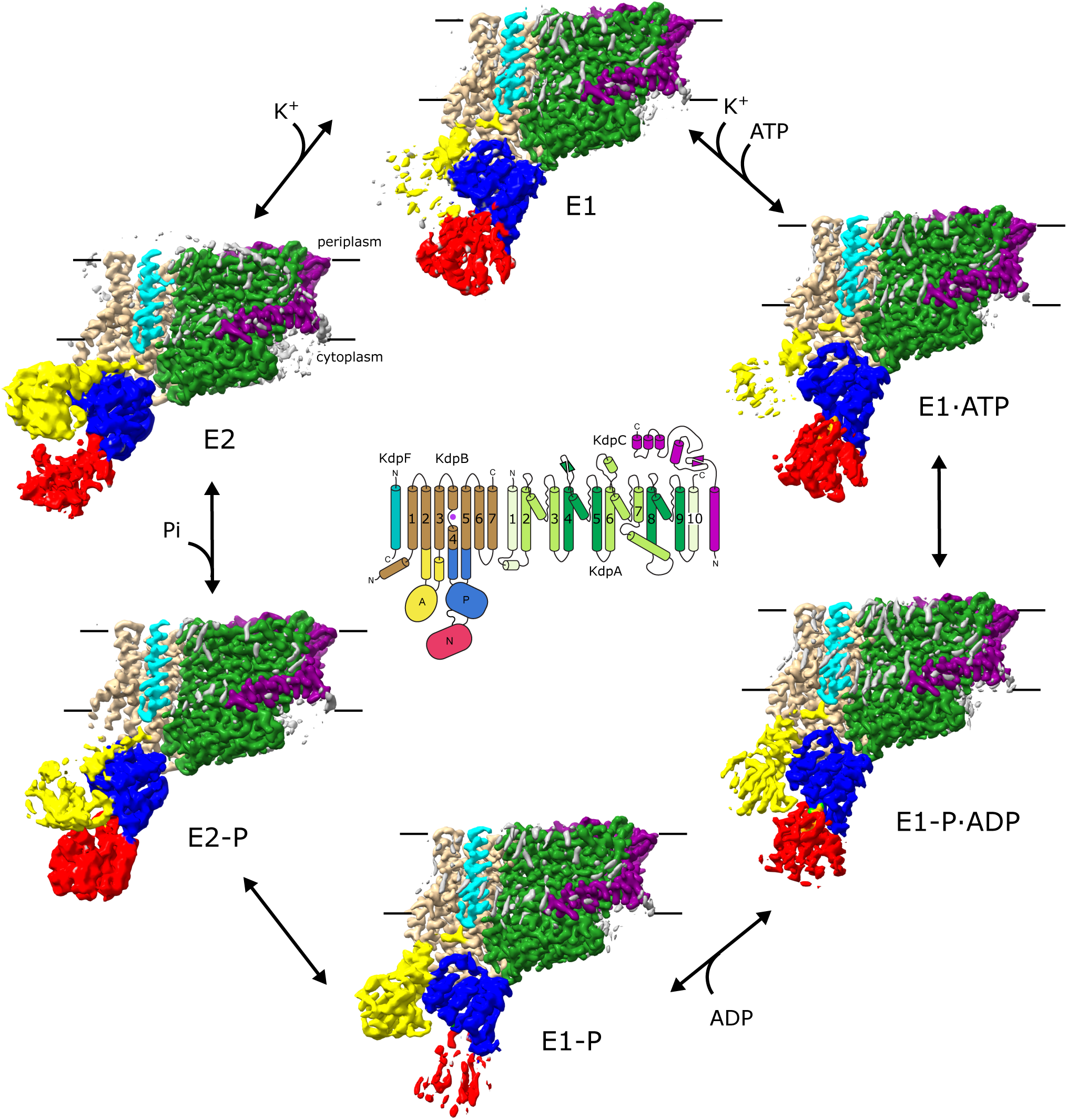
Cryo-EM structures of KdpFABC in lipid nanodiscs. Six density maps were derived from the images recorded under turnover conditions and assigned to a specific intermediate state in the Post-Albers reaction cycle (arrows). The diagram in the middle depicts helical topology and the coloring scheme that has been applied to the density maps. These density maps are hybrid structures with the membrane domains derived from conventional reconstruction with cryoSPARC and cytoplasmic domains with RELION. The membrane region is indicated by horizontal black lines.

**Figure 2.**
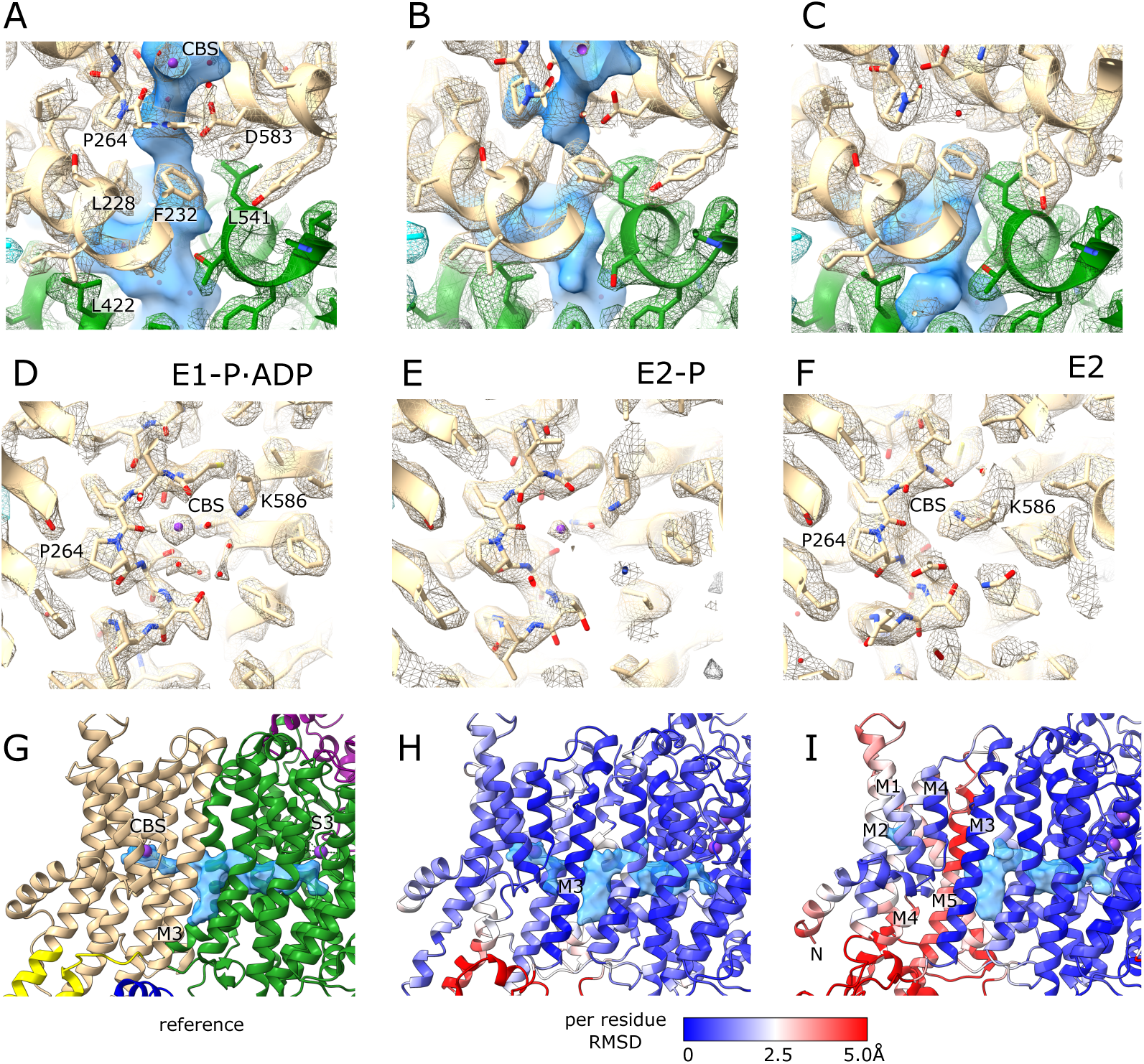
Structural dynamics of KdpFABC. (A-C) Structural changes at the KdpA/KdpB subunit interface cause tunnel occlusion as KdpFABC transitions from E1∼P through E2-P to E2. The tunnel is shown as a blue surface and map densities as a mesh with subunits colored as in Fig. 1. (D-F) Structural changes and K^+^ binding at the CBS during this transition. Although K^+^ remains bound and there is little structural change from E1∼P to E2-P, Lys586 swings into the CBS to displace the K^+^ in the E2 state. (G-I) Transmembrane domains of E2-P and E2 states are color coded to show per residue RMSD during the transition. The E1∼P·ADP state with standard coloring from Fig. 1 was the reference for RMSD calculation after aligning M3 helices from KdpB. Only modest movements accompany the E1∼P to E2-P transition, whereas the transition to E2 induces large movements including vertical displacement of KdpB-M5 by 4.2 Å (based on Cα of Lys586). Overall RMSD values are listed in Table 2.

**Table 1.** Summary statistics for structure determination.

| **Deposition** | **E1** | **E1-ATP** | **E1P** | **E2P** | **E2** |
| --- | --- | --- | --- | --- | --- |
| PDB | 9ZWK | 9ZWL | 9ZWM | 9ZWN | 9ZWO |
| EMDB | EMD-74910 | EMD-74911 | EMD-74912 | EMD-74913 | EMD-74914 |
| **Data collection and processing** |  |  |  |  |  |
| Magnification | 130kx | 130kx | 130kx | 130kx | 130kx |
| Voltage (kV) | 300 | 300 | 300 | 300 | 300 |
| Electron exposure (e<sup>-</sup>/Å<sup>2</sup>) | 50 | 50 | 50 | 50 | 50 |
| Defocus range (μm) | 1.0-3.0 | 1.0-3.0 | 1.0-3.0 | 1.0-3.0 | 1.0-3.0 |
| Pixel size (Å) | 0.93 | 0.93 | 0.93 | 0.93 | 0.93 |
| Symmetry imposed | C1 | C1 | C1 | C1 | C1 |
| Initial particle images (no.) | 10,382,000 | 10,382,000 | 10,382,000 | 10,382,000 | 10,382,000 |
| Final particle images (no.) | 89,556 | 270,688 | 273,670 | 141,132 | 78,696 |
| Map resolution* | 2.34/3.5 | 2.28/3.2 | 2.25/3.2 | 2.58/3.4 | 2.70/3.5 |
| FSC threshold (Å) | 0.143 | 0.143 | 0.143 | 0.143 | 0.143 |
| B factor (Å<sup>2</sup>) | 50.7/98.3 | 60.9/100.6 | 59.1/96.7 | 69.3/49.0 | 62.8/113.7 |
| **Model Refinement** |  |  |  |  |  |
| Model composition |  |  |  |  |  |
| Non-hydrogen atoms | 11017 | 11097 | 11067 | 10914 | 11000 |
| Protein residues | 1452 | 1455 | 1455 | 1445 | 1449 |
| Waters | 101 | 120 | 116 | 49 | 56 |
| Ligands | 5 | 6 | 6 | 5 | 5 |
| RMS deviations |  |  |  |  |  |
| Bond lengths (Å) | 0.002 | 0.002 | 0.002 | 0.002 | 0.002 |
| Bond angles (°) | 0.465 | 0.490 | 0.477 | 0.454 | 0.466 |
| Validation |  |  |  |  |  |
| MolProbity score | 1.57 | 1.35 | 1.47 | 1.73 | 1.51 |
| Clashscore | 6.79 | 4.88 | 5.43 | 6.64 | 6.99 |
| Rotamer outliers (%) | 0.70 | 1.32 | 1.14 | 2.65 | 0.97 |
| CaBLAM outliers (%) | 2.02 | 0.0 | 1.81 | 1.96 | 2.23 |
| Rama-Z score | 0.83 | 0.62 | 0.63 | 0.22 | -0.44 |
| Ramachandran plot |  |  |  |  |  |
| Favored (%) | 96.81 | 98.20 | 97.30 | 97.84 | 97.36 |
| Allowed (%) | 3.19 | 1.80 | 2.70 | 2.16 | 2.64 |
| Disallowed (%) | 0.00 | 0.00 | 0.00 | 0.00 | 0.00 |
| Qscore |  |  |  |  |  |
| Core TM domain | 0.80 | 0.81 | 0.82 | 0.71 | 0.73 |
| KdpB cytoplasmic domains | 0.25 | 0.37 | 0.28 | 0.37 | 0.22 |
| Model vs. Data CC (mask) | 0.84 | 0.86 | 0.86 | 0.78 | 0.80 |

\* The first value refers to the global refinement in cryosparc and the second value to the masked refinement of KdpB cytoplasmic domains in RELION.

**Table 2.** Structure comparisons relative to E1∼P·ADP*.

|  | E1 | E1·ATP | E1~P·ADP | E1~P | E2-P | E2 |
| --- | --- | --- | --- | --- | --- | --- |
| FABC subunits | 3.1<br>(1452) | 2.3<br>(1455) | - | 3.7<br>(1455) | 8.5<br>(1443) | 11.9<br>(1447) |
| A subunit | 0.2<br>(557) | 0.2<br>(557) | - | 0.2<br>(557) | 0.6<br>(557) | 0.6<br>(557) |
| B subunit | 4.6<br>(678) | 3.4<br>(681) | - | 5.4<br>(681) | 12.5<br>(671) | 17.4<br>(673) |
| B subunit TM | 0.5<br>(280) | 0.4<br>(283) | - | 0.4<br>(283) | 1.8<br>(273) | 3.2<br>(275) |
| C subunit | 0.3<br>(188) | 0.2<br>(188) | - | 0.2<br>(188) | 0.3<br>(186) | 0.3<br>(188) |
| F subunit | 0.3<br>(29) | 0.4<br>(29) | - | 0.3<br>(29) | 0.5<br>(29) | 0.7<br>(29) |

\* values represent RMSD in Å for Cα atoms. Number of atoms in parentheses.

Assignment of substates to intermediates in the Post-Albers transport cycle was determined by phosphorylation status of the conserved Asp307 at the catalytic site in the P-domain as well as the juxtaposition of the surrounding A- and N-domains. Like related P-type ATPases (17), the N-domain binds ATP and associates with the P-domain in E1 states in order to transfer γ-phosphate to Asp307. In contrast, the A-domain associates with the P-domain in E2 states in order to hydrolyze the resulting aspartyl phosphate. As reported previously (9), the class with the largest number of particles represents E1∼P·ADP; it features well-ordered cytoplasmic domains, the presence of Mg·ADP in the N-domain and phosphorylation of Asp307 in the P-domain (Sfig. 3). A second E1 state displays an ordered N-domain with density in the ATP site; however, Asp307 does not appear to be phosphorylated and the A-domain is disordered. These features are consistent with an E1·ATP state prior to phosphorylation. A third E1 state displays a disordered N-domain, density consistent with Asp307 phosphorylation and an ordered A domain, suggesting the E1∼P state after release of ADP and prior to transition to E2-P. The last E1 state features disordered A-and N-domains as well as a poorly ordered P-domain with no clear density in the vicinity of Asp307. In this last state, membrane helices of KdpB are congruent with other E1 states (RMSD of 0.4-0.5 for 280 Cα atoms, Table 2), so we have assigned it to apo E1 intermediately prior to binding ATP. All E1 states have strong density at the CBS which has been modeled as K^+^ (Sfig. 3).

**Figure 3.**
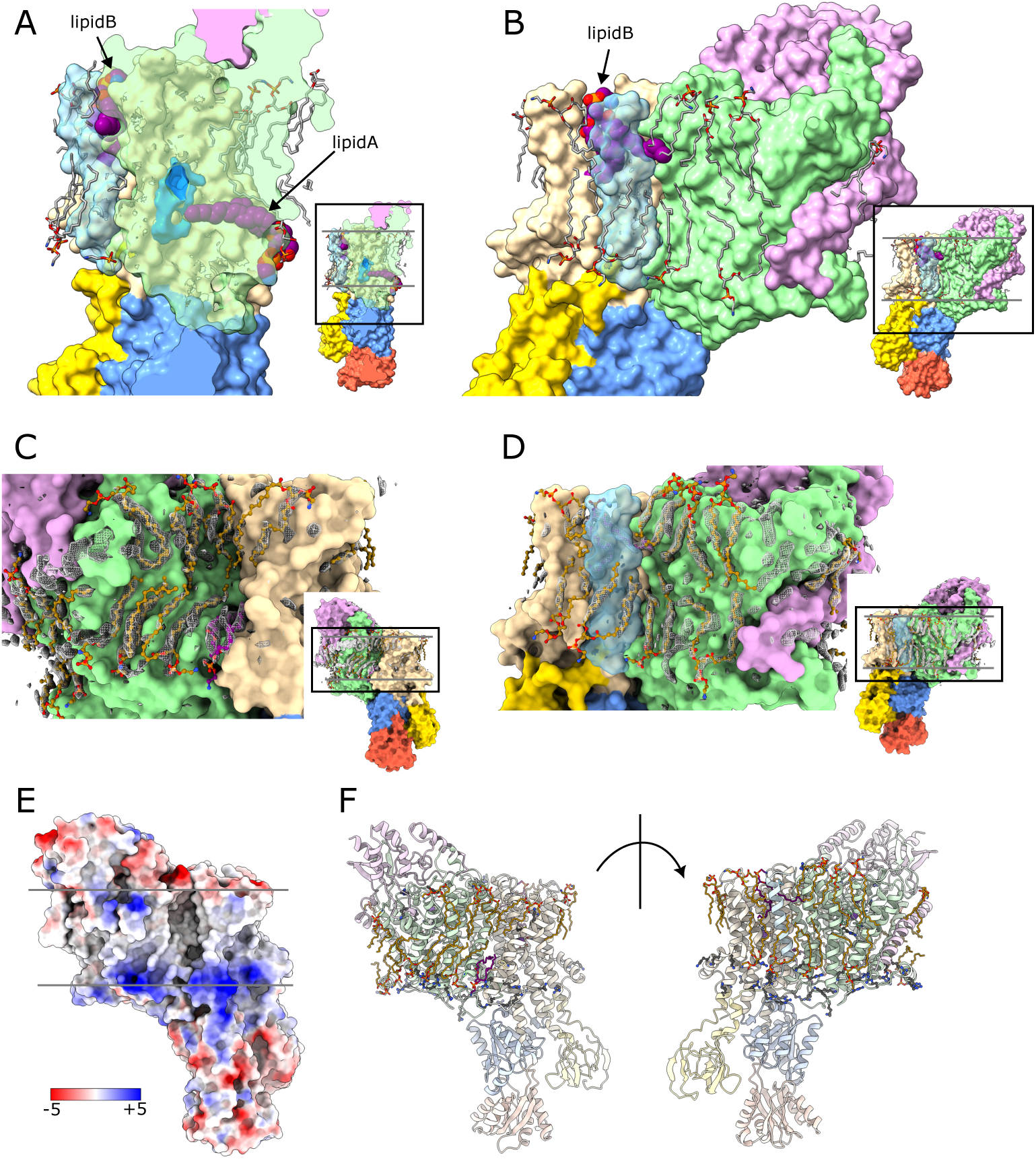
Lipid interactions with KdpFABC. (A-B) Two structural lipids bind at subunit interfaces. Lipid A binds at the cytoplasmic side of the membrane with its tail penetrating between KdpA and KpdB and contacting the intramembrane tunnel at its pinch point. Lipid B binds at the periplasmic side of the membrane with both tails wrapping around KdpF and mediating its interface with KdpB. KdpA and KdpF are rendered transparent in panels A and B, respectively. Inserts illustrate the area shown in the main panel (block rectangle) with grey horizontal lines indicating membrane boundaries. (C-D) Lipids were fitted to densities at the periphery of KdpFABC. The mesh was derived by aligning and summing the individual maps, though comparable densities were seen in all of the individual nanodisc structures (c.f. grey densities in Fig. 1). (E) The surface of KdpFABC is colored according to coulombic potential, showing a prevalence of positive charge at the cytoplasmic membrane surface. (F) The prevalence of arginine and lysine residues, shown in stick representation, account for this positive charge. Annular lipids are shown in gold, whereas lipids A and B are purple.

Our structures include two E2 states, which are distinguished from E1 states by large movements of the cytoplasmic domains. The first has a moderately well ordered A-domain in which the conserved TGES^162^ loop is adjacent to a phosphorylated Asp307 residue (Sfig. 3), making it consistent with E2-P. The membrane domain of KdpB has undergone modest changes (RMSD relative to E1∼P·ADP of 1.8 Å for 273 Cα atoms, Table 2), though the helices have maintained their register and a spherical density, modeled as K^+^, is still visible at the CBS (Sfig. 4). The second E2 state has bigger differences including an ∼4-Å translation of the M5 helix of KdpB and absence of density at the CBS (Sfig. 4); RMSD for the membrane domain of KdpB is 3.2 Å (275 C<u>α</u> atoms, Table 2). Additionally, Asp307 does not appear to be phosphorylated and the A-domain has moved away from this catalytic center (Sfig. 3), consistent with the post-hydrolysis E2 state.

**Figure 4.**
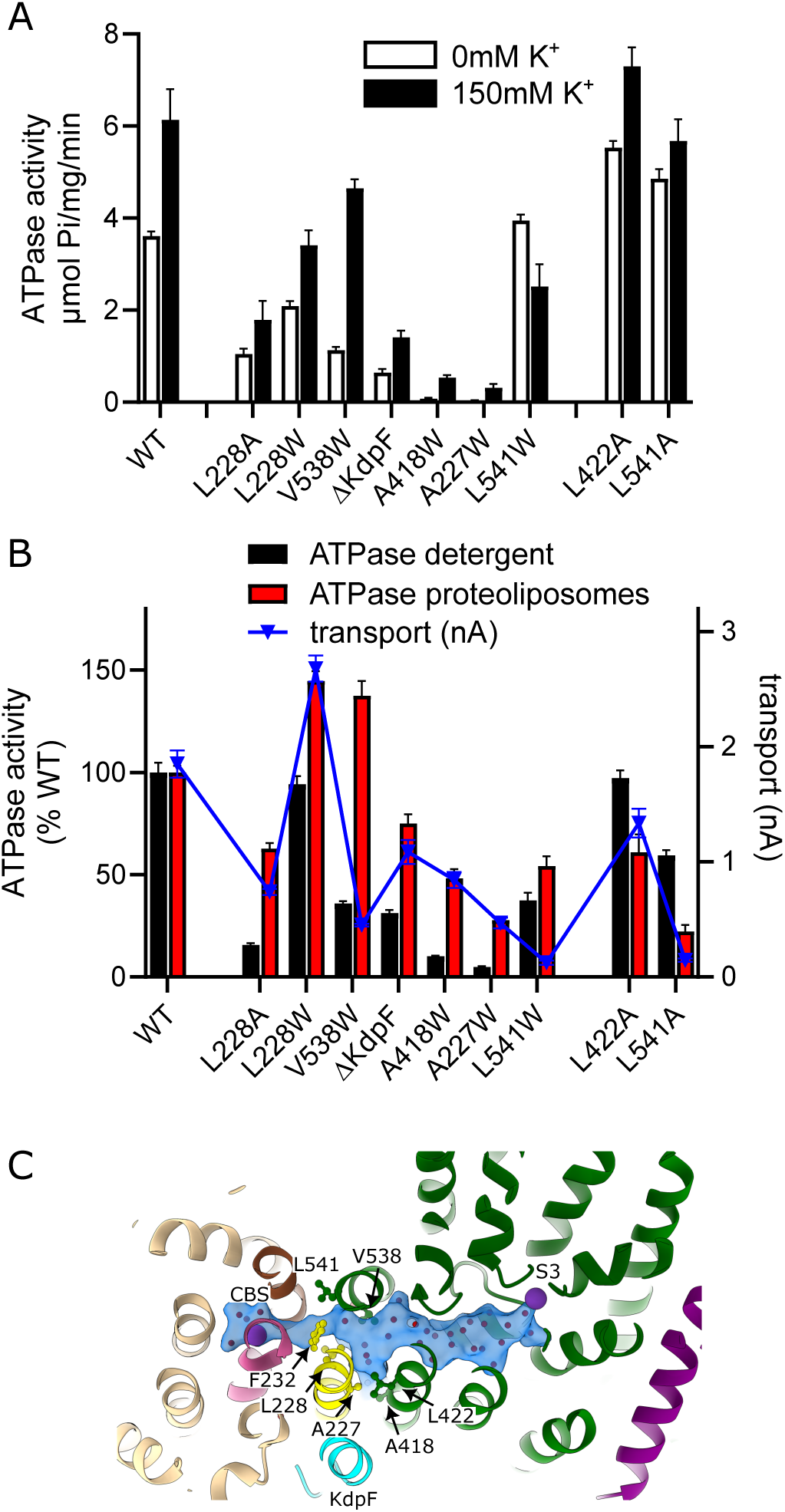
ATPase and transport activity of KdpFABC mutants. (A) Mutations have variable effects on ATPase activity of detergent solubilized samples. Like WT, all mutants except L541W show stimulation upon addition of K^+^. (B) Comparison of ATPase activity of proteoliposome reconstituted (red bars) and detergent-solubilized (black bars) samples, both of which have been normalized relative to WT (left axis). Transport activity of proteoliposome samples (blue triangles, right axis) reflect energy coupling. ATPase and transport activities for proteoliposome data were compensated for reconstitution efficiency as quantified by SDS PAGE (Sfig. 7). (C) Location of mutants, which are all near the KdpA/KdpB interface; Leu228, Ala227 are on M3 of KdpB (yellow) and Ala418, Leu422, Val538 and Leu541 are on M9 and M10 of KdpA. M4 and M5 of KdpB are colored pink and dark brown, respectively (also in Fig. 6 and Movie 1 & 2). Error bars represent SEM based on 3-6 technical replicates.

KdpA, KdpC and KdpF are well resolved in all the maps with resolutions <2.5 Å (Sfig. 1). These subunits are essentially static throughout the transport cycle, with RMSD <0.7 Å for all states (Table 2). In particular, the selectivity filter is well resolved with very strong density at the S3 position (Sfig. 5), consistent with its role as the primary site for K^+^ binding and selectivity (9). In the five new structures, strong density is also seen at the S1 site, which has therefore been modeled with a second K^+^ at the mouth of the selectivity filter. In contrast to a previous cryo-EM analysis under turnover conditions (6), we did not detect the auto-inhibited E1∼P conformation. This discrepancy is due to genetic knockout of KdpD in our expression strain, which is responsible for post-translational phosphorylation of Ser162 in KdpB and generation of this inhibited conformation (18, 19).

**Figure 5.**
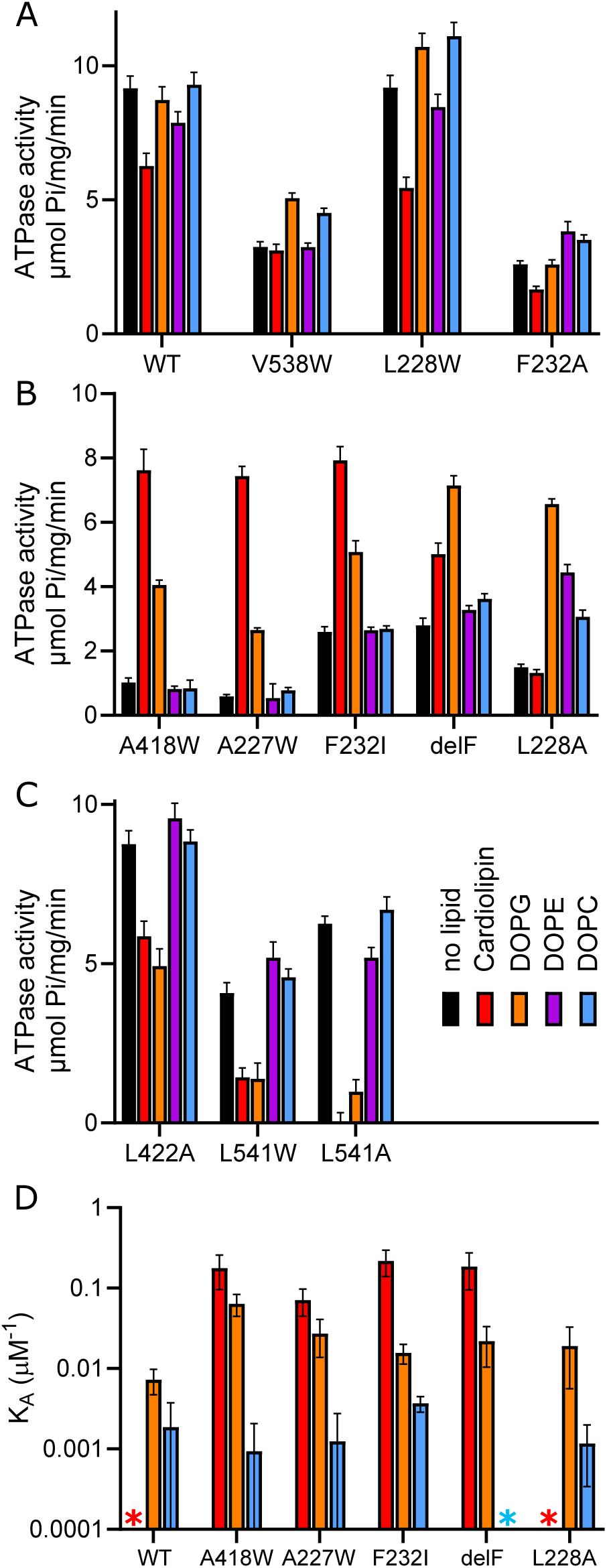
Lipid effects on ATPase activity. (A-C) Effects of two negatively charged (CL and DOPG) and two neutral (DOPE and DOPC) lipids on ATPase activity of detergent-solubilized preparations. Full titrations of a wider range of lipids are shown in Sfig. 8. Panel A includes WT and three mutants with relatively little change in activity after addition of lipids. Panel B includes five mutants showing marked stimulation by negatively charged lipids, whereas neutral lipids had little to no effect. Panel C includes three mutants that were inhibited by negatively charged lipids but unaffected by neutral lipids. Error bars in A-C represent SEM based on fit to corresponding titrations in Sfig. 8, which include triplicate measurements at six lipid concentrations. (D) Binding affinity of mutants from panel B was measured by MST, showing higher affinity binding of negatively charged lipids. Astericks indicate measurements with insufficient change in MST signal, indicating very low affinity. Full MST titrations are shown in Sfig. 9; error bars in panel D represent SEM based on fit to these titrations, which include tripilicate measurements of 14 concentrations.

### Sequence of key structural changes driving the transport cycle

Assignment of structures relative to intermediate states of the transport cycle together with increased resolution relative to previous work allows us to better define the sequence for two key structural changes. The first involves occlusion of the intramembrane tunnel, which is essential for preventing backflow of K^+^ and thus ensuring that ion transport is energetically coupled to ATP hydrolysis (7). Increased precision of side chain rotamers affords a more reliable definition of the tunnel profile and structural changes associated with occlusion. To aid this definition, we adopted a plugin to pyMol called Cavitomix (20). Unlike Caver (21) used in previous work (8, 9), Cavitomix generates a non-spherical cross-section of internal cavities that is a more realistic rendering of this intramembrane tunnel. Consistent with previous reports, the tunnel extends from the selectivity filter of KdpA to the CBS of KdpB in all the E1 states (Sfig. 6), thus allowing K^+^ to transit between its binding sites. As the complex transitions from E1∼P to E2-P, the tunnel is pinched near the KdpA/KdpB interface and then completely collapses in the E2 state (Fig. 2, Movies 1 & 2). The pinch point is surrounded by Val538 and Leu541 on KdpA as well as Leu228 and Phe232 on KdpB; in addition, the aliphatic tail of a lipid molecule penetrates the subunit interface and seals the tunnel. In previous work, Phe232 was shown to be essential for occlusion (5); however Phe232 does not move during the transition. Instead, the two leucines and the lipid tail encroach on the tunnel in the E2-P state, thus accounting for a narrower profile that would block passage of water and ions. This pinching is significantly enhanced in the E2 state, in which the cavity completely collapses in KdpB.

**Figure 6.**
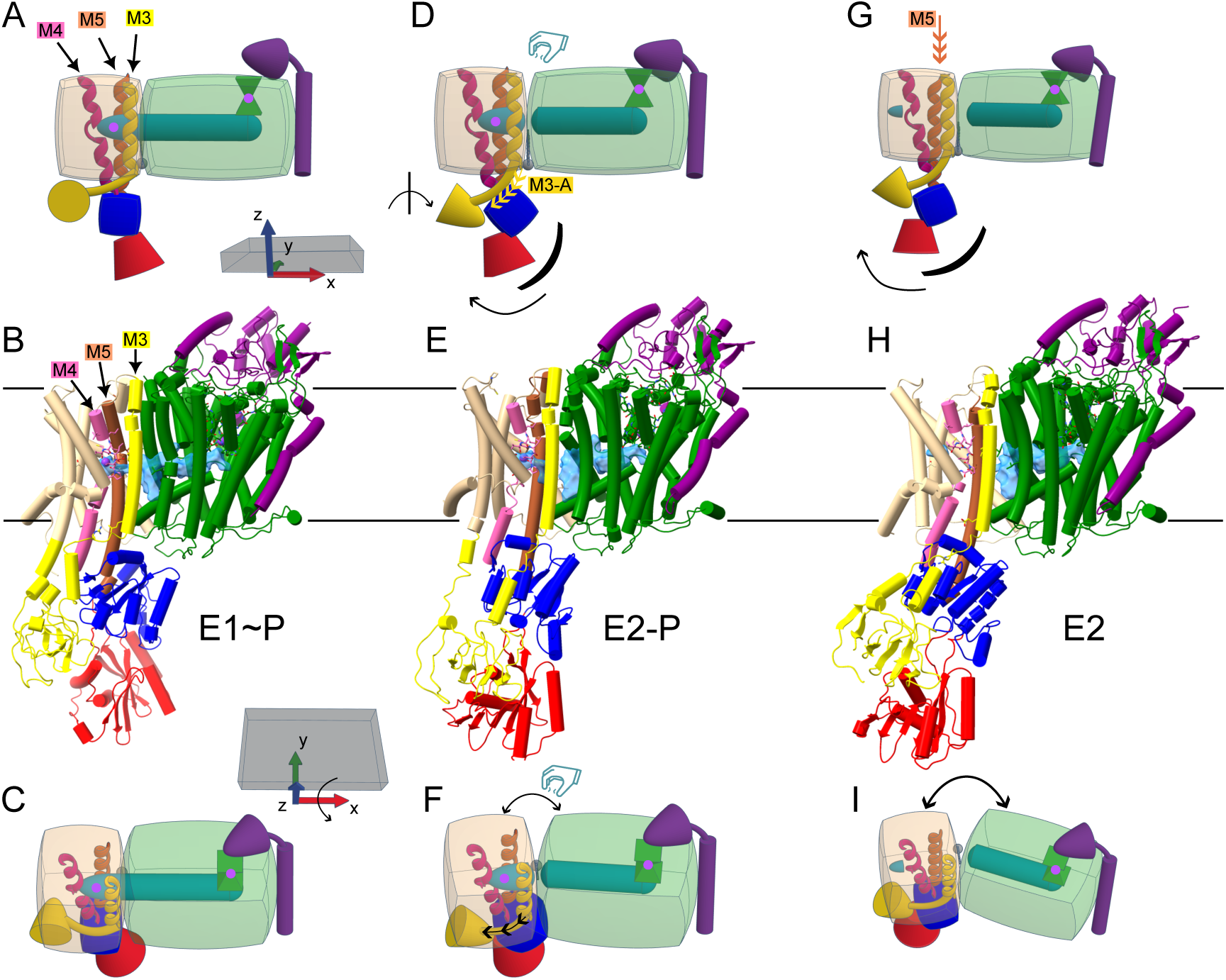
Allosteric coupling of KdpFABC. Diagrams illustrating key features of the structural dynamics in the transport cycle. E1∼P (A-C), E2-P (D-F) and E2 (G-I) are the same states shown in Fig. 2. The E1∼P to E2-P step generates ∼90° rotation of the A-domain and inclination of N- and P-domains. A-domain rotation strains its linker with M3 which leads to rotation of KdpA relative to KdpB, thus pinching off the intramembrane tunnel (blue). The E2-P to E2 step leads to further rotation of KdpA and inclination of N- and P-domains and displacement of the M5 helix perpendicular to the membrane plane. Movement of M5 places the side chain of Lys586 in the CBS (Fig. 2) and, together with movement of other KdpB elements, collapses the cavity surrounding the CBS. Instead, a small cavity opens to the left of M4, constituting the exit site for K^+^, as previously described (9). The top two rows represent views parallel to the membrane plane (x-y) whereas the bottom row has been rotated ∼70° around the x axis. KdpF was removed for clarity and colors are as defined in Fig. 1, except that M3, M4 and M5 and KdpB are colored yellow, pink and brown, respectively.

The second key structural change displaces K^+^ from the CBS and can be thought of as the “power stroke” of the reaction cycle. In other P-type ATPases, the CBS typically transitions from high to low affinity during the pumping cycle (17). For KdpB, however, recent evidence suggests that Lys586 pushes the ion into a low affinity exit site located near Thr75 on M2 (8, 9). Our new structures show unambiguous density for Lys586 in both E2 states and confirms swinging of its side chain into the CBS during the E2-P to E2 transition as M5 translates >4 Å toward the cytoplasm (Fig. 2f,i and Sfig. 4). The E2-P state features a K^+^ at the CBS next to a water-filled cavity. In the E2 state, however, the CBS is empty and, as mentioned above, this cavity collapses. Instead, a new cavity with a spherical density (likely K^+^ but modeled as water) appears next to Thr75 (Fig. 2f, Sfig. 6), consistent with its hypothesized role as the low-affinity release site for K^+^ (9).

More general comparison of the membrane helices show that large scale changes occur during this transition from E2-P to E2. Specific movements are illustrated with per-residue deviations in Fig. 2h-i (RMSD of Cα atoms after alignment of M3 from the E1∼P·ADP state). This analysis reveals relatively small movements during the E1∼P to E2-P transition and much larger movements in the subsequent transition to E2 (also see Table 2 and Movies 1 & 2). Although most KdpB helices are dynamic, the largest movements are in M5 and the cytoplasmic half of M4 - distal to Pro264 and the CBS. Together, these movements indicate that the power stroke in K^+^ transport occurs upon hydrolysis of the aspartyl phosphate to produce the E2 state.

### Lipids bind to the KdpFABC complex

Previous structures in detergent solution (4–6, 8) feature elongated, non-protein densities at the periphery of the complex, which are generally too noisy to assign to detergent or lipid. The current structures from nanodisc-reconstituted protein reveal similar densities, though with greater consistency at a larger number of sites (c.f. grey, elongated densities in Fig. 1). Given the absence of detergent in this sample, they have been modeled as lipids. Two particularly well-resolved lipids are present in all our structures and penetrate deeply into the complex at subunit interfaces. The first, dubbed lipid A, has two well-defined aliphatic tails intercalating between KdpA and KdpB; one tail lines the tunnel as it passes its pinch point at the interface of these subunits (Fig. 3a). Lipid A has been modeled in previous structures, though the weak density for the headgroup make its chemical nature uncertain. As in our previous structures, we have modeled it as phosphatidylethanolamine (PE) as suggested by earlier mass spectroscopy results (8), though Silberberg et al. (5) modeled it as cardiolipin based on results from coarse-grain MD simulations. The second density, dubbed lipid B, resides at the periplasmic side of the complex with its headgroup near the N-terminus of KdpF (Fig. 3b). Two aliphatic tails from lipid B are well resolved in all our structures wrapping around KdpF and thus mediating its interaction with KdpB.

In addition to these well-ordered interface lipids, additional densities are visible around the periphery of the complex, forming a belt within the plane of the membrane. These likely represent annular lipids and they occupy consistent positions in all our maps, though individual densities are difficult to model due to their discontinuity (Fig. 1). To elucidate the distribution of these annular lipids more clearly, we produced a consensus map by aligning the individual maps from all six substates and, after subtracting the portion of the map corresponding to protein, summing the remaining densities. Figs. 3c and d show this consensus map with the peripheral densities modeled as 20 dioleoyl phosphatidylethanolamine (DOPE) molecules, though head group assignment is arbitrary given a general lack of map density.

### Perturbations to the tunnel and subunit interface

Structural dynamics of the intramembrane tunnel (Fig. 2), documented effects of KdpB-Phe232 (5), and a large energy barrier detected by MD simulation (15) all point to the KdpA/KdpB interface as key to understanding coupling of K^+^ transport to ATP hydrolysis. To study this region further, we introduced mutations to perturb the interface and/or interfere with tunnel constriction. Specifically, mutations of Ala418, Leu422, Val538, Leu541 in KdpA and Ala227, Leu228 in KdpB were generated and initially evaluated by measuring ATPase activity in detergent solution (Fig. 4a). For WT protein, ATPase is dependent on addition of K^+^, as expected for a well coupled K^+^ pump. Given the high K^+^ affinity of KdpFABC, this measurement requires reduced concentrations of Mg·ATP to minimize K^+^ contamination from this reagent (9). Under these conditions, ATPase activity of WT is stimulated ∼1.6 fold by addition of 150 mM K^+^. With the exception of L541W, all of the mutants displayed similar K^+^ stimulation, indicating that the pumping unit in KdpB remained allosterically coupled. However, it is notable that maximal ATPase levels were reduced to a varying extent for most of the mutants.

We went on to assess transport efficiency and energy coupling by reconstituting mutant complexes into liposomes and comparing levels of ATP hydrolysis with associated membrane current. Given potential differences in reconstitution efficiency, gel electrophoresis was used to quantify protein incorporation into liposomes and then to normalize activities relative to WT (Sfig. 7). With respect to ATPase activity, comparison of detergent solubilized and reconstituted preparations indicated that several mutants (A418W, A227W, L228A, L228W, V538W) are stimulated by the lipid environment, whereas others (L541A, L422A) are inhibited (Fig. 4b, red vs. black bars). With respect to energy coupling, comparison of transport and ATPase activities showed that several mutants were compromised, i.e., with diminished transport relative to their respective ATPase activities (Fig. 4b, red bars vs. blue triangles for V538W, L541W and L541A). Such uncoupling of transport from ATP hydrolysis suggests that the corresponding mutation interferes with occlusion of the tunnel, thus allowing K^+^ to escape back into the periplasm during the power stroke.

In addition to these mutants, we investigated deletion of KdpF, which comprises a single-pass membrane helix that is not present in all bacterial species (3). Previous work showed that deletion of KdpF from the *E. coli* complex resulted in almost complete inactivation of ATPase in amine oxide detergent, but that activity was recovered by addition of lipid (16). In our hands, the ΔKdpF complex had ∼20% of WT activity in decyl-maltoside (DM), which is a milder detergent (Fig. 4a). After reconstitution of the ΔKdpF construct into proteoliposomes, ATPase was increased ∼2.5 fold and produced transport currents consistent with a well coupled complex (Fig. 4b), confirming the lipid sensitivity of this complex.

### Negatively charged lipids stabilize the KdpFABC complex

To further investigate the influence of lipids, we measured ATPase activity of detergent-solubilized preparations supplemented by a wide variety of lipid species. For these assays, we tested a variety of head groups with different length and saturation of aliphatic tails. Although addition of lipid to WT protein produced only minor effects, a number of mutants had dramatic responses (Fig. 5, Sfig. 8). The largest effects were observed with KdpB-A227W and KdpA-A418W (Fig. 5b), which had very low activity in the absence of lipid, but were stimulated 4- to 12-fold by the addition of negatively charged lipids, namely cardiolipin (net −2), dioleoyl phosphatidylglycerol (DOPG) and DOPA (both net −1). A227W had a preference for cardiolipin, whereas A418W was stimulated by a broader selection of negatively charged lipids that included both saturated and unsaturated tails (Sfig. 8). The opposite effect was observed for KdpA-L541A/W and KdpA-L422A mutants, for which negatively charged lipids had an inhibitory effect. Consistent with previous work, (16), ΔKdpF was strongly lipid dependent; current work shows that this effect is once again attributable to negatively charged lipids. Interestingly, KdpB-F232I was strongly stimulated by negative lipids, but KdpB-F232A was not. Mutation of F232 was previously shown to not only uncouple ATPase from transport, but also to eliminate K^+^ dependence (5, 15).

Microscale thermophoresis was used to document binding of lipids to detergent-solubilized KdpFABC mutants (Fig 5d, Sfig. 9). WT KdpFABC showed no discernable interaction with cardiolipin, and relatively low affinity binding of DOPG and DOPC (Ka < 0.01 μM^-1^). In contrast, A418W, A227W, A228A, F232I and ΔKdpF display 10-100 fold higher affinity interaction with negatively charged species relative to DOPC. Furthermore, these binding data were largely consistent with effects observed on ATPase activity.

Finally, we used mass spectrometry to document the lipid species associated with detergent-solubilized, WT KdpFABC. After size-exclusion chromatography of affinity-purified protein in DM detergent, PE and PG lipids were most abundant, comprising ∼43% and ∼31% of the total with the remainder comprising a variety of cardiolipins (Table 3). To identify the more tightly bound lipids, we evaluated these same samples after reconstitution into nanodiscs. Reconstitution involves mixing with additional detergent and exogenous lipids followed by another purification by size-exclusion chromatography, thus causing depletion of weakly bound lipids. To compare samples before and after reconstitution, we excluded the exogenous lipids PC and PA during processing of the spectral data. We found four PG species and one PE species to be enriched after nanodisc reconstitution (Table 3). In contrast, all cardiolipin species were substantially reduced. We conclude that PG and PE co-purify with WT KdpFABC and are therefore more tightly bound than cardiolipin.

**Table 3.** Lipid analysis by mass spectrometry.

| Lipid species | Abundance<br>Detergent<br>(per 100<br>molecules) | Abundance<br>Nanodisc<br>(per 100<br>molecules) | Log<sub>2</sub>(Nano/Det) |
| --- | --- | --- | --- |
| Cardiolipin |  |  |  |
| CL(16:0_16:0_16:0_16:1) | 1.142 | 0.274 | -2.0589 |
| CL(14:0_16:0_16:1_18:1) | 4.496 | 0.667 | -2.7532 |
| CL(16:0_16:0_16:1_17:1) | 2.849 | 0.505 | -2.49494 |
| CL(16:0_16:0_16:0_18:1) | 0.558 | 0.125 | -2.15945 |
| CL(16:0_16:0_16:1_18:1) | 5.662 | 1.434 | -1.98125 |
| CL(16:0_16:1_16:1_18:1) | 1.447 | 0.365 | -1.9853 |
| CL(16:0_16:0_17:1_18:1) | 2.658 | 0.902 | -1.55869 |
| CL(16:0_16:0_18:1_18:1) | 3.704 | 1.531 | -1.27477 |
| CL(16:0_16:1_18:1_18:1) | 1.578 | 0.841 | -0.90796 |
| CL(16:0_16:0_18:1_19:1) | 1.257 | 0.288 | -2.12739 |
| PG |  |  |  |
| PG(16:0_16:0) | 1.818 | 0.387 | -2.23365 |
| PG(16:0_16:1) | **14.010** | **18.354** | **0.389657** |
| PG(16:0_17:1) | 9.428 | 5.475 | -0.78418 |
| PG(16:0_18:1) | **15.148** | **24.418** | **0.688797** |
| PG(16:1_18:1) | **2.306** | **16.518** | **2.840509** |
| PG(17:1_18:1) | **0.793** | **2.519** | **1.667903** |
| PE |  |  |  |
| PE(16:0_18:1) | 23.828 | 8.153 | -1.54715 |
| PE(16:1_18:1) | **7.318** | **17.244** | **1.236553** |

## Discussion

In this work, we have used cryo-EM to image KdpFABC embedded in a lipid bilayer under active turnover conditions. The resulting structures represent a comprehensive description of the transport cycle and the improved resolution sheds new light on two key events: occlusion of the intramembrane tunnel that carries K^+^ through the complex, and subsequent displacement of this ion into the low-affinity release site. These structures also reveal binding sites for ∼20 lipid molecules. The majority are annular lipids residing around the periphery of the complex, but two lipids with well-ordered tails are bound at distinct subunit interfaces. A functional role of negatively charged lipids has been uncovered by introducing mutations at the KdpA/KdpB interface or by deleting the KdpF subunit.

### Lipid-protein interactions

A large body of work has addressed interactions between lipids and a diverse array of membrane proteins, which has been reviewed extensively (22, 23). Lipids can be divided into three categories: bulk lipids, annular lipids and structural lipids. Bulk lipids constitute the matrix of the membrane; although they do not directly contact the embedded proteins, bulk lipids can nevertheless regulate function by influencing physical-chemical properties of the bilayer such as fluidity, thickness and lateral pressure. Annular lipids directly contact the perimeter of the protein, interacting either specifically or non-specifically via hydrophilic head group and/or aliphatic tails. Annular lipids, which exchange readily with the bulk, are thought to be important in reconciling hydrophobic mismatch and in facilitating conformational change. Finally, structural lipids play a wide variety of roles in mediating protein interactions and filling fenestrations in the protein surface.

We found two structural lipids bound to KdpFABC. Both lipids penetrate deeply into subunit interfaces and thus appear to be an integral part of this protein complex. Lipid A, which abuts the intersubunit tunnel, has in fact been observed in all previous cryo-EM structures of KdpFABC prepared either in decyl-maltoside (8) or in dodecyl-maltoside (5–7) detergents. Although not modeled in the X-ray structure from KdpFABC prepared in octyl-glucoside (4), density for lipid A is visible in one of three copies composing the unit cell. Given that lipid A seals a large gap in the protein at the site of tunnel constriction, it seems likely that delipidation would compromise the continuity of this K^+^ conduit.

The second structural lipid - lipid B - resides between KdpB and KdpF on the periplasmic side of the membrane. Although lipid B has not previously been described, its density is visible in higher resolution cryo-EM maps (EMD-12184, EMD-23268, EMD-14917 at 2.9-3.2 Å resolution) from both decyl-maltoside and dodecyl-maltoside solubilized preparations (6, 8). Both tails of lipid B intercalate between KdpF and M3/M4 helices of KdpB such that the periplasmic end of KdpF does not make direct contact with KdpB. Like previous work (16), deletion of KdpF severely compromises ATPase activity in detergent solution, but activity is recovered by addition of exogenous lipids and transport is observed after reconstitution into proteoliposomes. These data imply that the ΔKdpF complex is readily delipidated and that lipid B somehow facilitates the conformational changes in KdpB that result in transport.

Our structures in lipid nanodiscs also revealed what appear to be annular lipids around the periphery of the complex. Despite controversy about the distinction between annular and bulk lipids (24), densities are clearly visible at consistent sites in the individual maps, suggesting somewhat stable binding sites on the surface of the protein. Most of these annular lipids are associated with KdpA and KdpC, presumably because these subunits remain static during the cycle. In contrast, KdpB undergoes major structural changes which likely results in more dynamic interactions with the surrounding lipids.

### Structural perturbations induce delipidation

Unlike previous work (5), addition of lipid did not have much effect on activity of detergent-solubilized WT protein. However, when Trp or Ala mutations were introduced to destabilize the subunit interface and interfere with the tunnel, we observed a variety of lipid-dependent behaviors. These include (1) stimulation of ATPase activity (A227W, L228A, L228W, A418W, V538W), (2) inhibition of ATPase activity (L422A, L541A) and (3) uncoupling of ATPase activity from transport (V538W, L541W, L541A). We propose that, similar deletion of KdpF, the first set of mutations rendered the complex prone to delipidation. Thus, the loss of activity in these mutants in detergent solution unmasks a functional role of lipid that is otherwise tightly bound by the WT complex. For the second set of mutations, the inhibitory effect of lipids might indicate that lipids are able to occupy sites that block the tunnel (e.g., deeper insertion of lipid A). Finally, for the third set of mutations with an uncoupling phenotype, we tentatively propose that the bulky side chain from the Trp substitution could either directly or indirectly block passage of K^+^ through the tunnel. Despite the diversity of effects, these results add further evidence for the subunit interface as a critical region for KdpFABC function.

Negatively charged lipids are primarily responsible for the observed functional effects on these mutants. In previous work, course-grain MD simulations concluded that cardiolipin, which carries two negative charges, preferentially associated with the complex (5) and this same study showed that cardiolipin stimulated ATPase activity of WT protein in dodecyl-maltoside detergent. We also observed that cardiolipin was potent in either stimulating or inhibiting ATPase activity of mutant complexes. However, cardiolipin is known to interact with a variety of proteins in vitro even in the absence of a defined physiological role (25). In the case of KdpFABC, our mass spectrometry results indicate that cardiolipin is not preferentially bound to WT protein and suggests that PG is the most likely species responsible for functional integrity of the complex.

As is common with membrane proteins (26), KdpFABC has a preponderance of positively charged residues (Arg and Lys) at the cytoplasmic membrane surface (Fig. 3e,f). However, given the lack of density for lipid head groups in our cryo-EM maps, a simple electrostatic interaction may not be the primary determinant of binding. On the other hand, the aliphatic tails are well resolved and display excellent shape complementarity with the protein surface, presumably reflecting strong van der Waals interactions. The importance of these interactions has been demonstrated in structural studies of aquaporin showing that lipids of varying head group composition occupy the same position on the protein surface (27). This result is consistent with the low specificity of annular lipids in general, which typically resemble the composition of the bulk membrane (28). Nevertheless, the overall positive surface potential could play a role in attracting negative lipids to the protein surface and thus explain their preferential effects on KdpFABC mutants.

### Allosteric coupling in transport

Our structures shed new light on two key steps responsible for coupling ATP hydrolysis to K^+^ transport. The first, E1∼P to E2-P, converts the high-energy aspartyl phosphate to a lower energy configuration. This step is thought to consume most of the energy derived from ATP (29) and, for KdpFABC, the prevalence of E1∼P species under turnover conditions (6, 9) suggests that it is rate-limiting. For conventional P-type ATPases, the E1∼P to E2-P step converts the CBS from high to low affinity and exposes this site to the extracellular side of the membrane in preparation for ion release. For KdpFABC, however, the CBS does not undergo the dramatic changes seen in other family members. Instead, this step is associated with occlusion of the intramembrane tunnel, leading to trapping of the K^+^ ion within the CBS.

Although closure of the tunnel was apparent from initial comparison of E1 and E2 states (7), the allosteric elements coupling movement of KdpB cytoplasmic domains to the tunnel have not been described. Current work indicates that tunnel occlusion is controlled by Val231, Phe232, Leu228 on M3 of KdpB as well as Ile422, Val538 and Ile541 on M8 and M10 of KdpA. Lipid A also contacts the tunnel at the pinch point and although the end of the aliphatic tail moves 2-3 Å during occlusion, it is likely to be in response to conformational change at the KdpA/KdpB interface. As documented in Table 2, there are negligible structural changes in KdpA, suggesting that the closure is driven mainly by KdpB. Like other P-type ATPases (29), the A-domain undergoes a large movement during the E1∼P to E2-P step. This movement is accompanied by tilting of the P-domain toward the membrane to enable the conserved TGES motif (specifically Glu161 in KdpB) to interact with the aspartyl phosphate (Asp307) in preparation for its hydrolysis. To accommodate movement of the A-domain, its linkage to M3 (Glu208 to Thr215) stretches out, thus breaking a salt bridge between conserved residues Arg212 and Glu218; this salt bridge is otherwise intact in all of our ligand-bound E1 structures (E1·ATP, E1∼P·ADP, E1∼P). In addition to these cytoplasmic domain movements, the E1∼P to E2-P step initiates rotation of the membrane domain of KdpB relative to KdpA about an axis roughly aligned with M3 (Movie 1). This movement coincides with subtle adjustments in Leu422 (M8) and Ile541 (M10) on KdpA, which are in van der Waals contact with M3 and M5 on KdpB. Although the causality of these movements is not clear, it seems plausible that breakage of the Arg212-Glu218 salt bridge may alleviate tension in the M3 helix, thus allowing the structure to relax into this occluded state.

The second key step in K^+^ transport by KdpFABC involves hydrolysis and release of the phosphate: E2-P to E2. A conformational change in KdpB during this step results in Lys586 swinging into the CBS and thereby displacing the K^+^ ion into the release site between M2, M4 and M6 (Fig. 2). This displacement was previously associated with the E1∼P to E2-P step (5, 8), but our structures clarify the timing of this key event, akin to the power stroke of molecular motors (30). The associated conformational change features a 4-5 Å, piston-like movement of the entire M5 helix on KdpB, which follows a further tilting of all three cytoplasmic domains toward the membrane surface (Movie 2). Additional movements include an unwinding of the M2/A-domain linker, which docks against the P-domain, and a lifting of the N-terminal, amphipathic helix that lies on the cytoplasmic membrane surface. The movements of M2, M5 and this N-terminal helix are consistent with deformation of the membrane in this region, as has been documented with SERCA, suggesting that potential energy may be stored in the membrane and somehow contribute to the allosteric coupling during this step (31). There may also be a role for annular lipids in alleviating hydrophobic mismatch caused by movements of these membrane helices out of the membrane plane (32).

### Summary

We have imaged fully active, WT KdpFABC in lipid nanodiscs under turnover conditions to generate high resolution structures representing all major intermediates of the transport cycle. Observed conformations are largely consistent with previous structures imaged in detergent micelles, confirming that KdpB undergoes dynamic changes in conjunction with ATP hydrolysis whereas KdpF, KdpA and KdpC remain almost completely static. Increased resolution of our structures sheds light on two key steps involving (1) occlusion of K^+^ at the CBS, due to closure of the intramembrane tunnel, and (2) active displacement of K^+^ from the CBS into the release site, which we view as the power stroke of the cycle. In addition, the structures reveal annular lipids around the periphery of the complex as well as two constitutively bound lipid molecules that appear to play an important structural role. Indeed, effects of delipidation were documented after introducing mutations at the interface of KdpA and KdpB or deletion of KdpF.

Finally, we propose that the linker between M3 and the A-domain of KdpB provides allosteric coupling responsible for coordinating ATP hydrolysis with movement of K^+^ through the unique intramembrane tunnel connecting entry and exit sites for this ion in KdpA and KdpB, respectively.

## Materials and Methods

### Cryo-EM data processing

Procedures for expressing and purifying WT and mutant KdpFABC complexes and for reconstitution into lipid nanodiscs have previously been described in detail (9, 33). For cryo-EM analysis, nanodisc-reconstituted WT KdpFABC was imaged after adding Mg·ATP to generate turnover; a large dataset of ∼50,000 images was collected and initial processing led to the previously reported structure of the E1∼P·ADP conformation (9). For the current work, this same dataset was further analyzed to generate five additional conformational states, as outlined in Sfigs. 1 and 2. Images were divided into several subsets for initial processing using cryoSPARC v4.6 (Structura Biotechnology Inc.), which after successive rounds of heterogenous refinement, culminated in three main particle classes, which are shown at the top of Sfig. 1. The corresponding particle sets were subjected to masked 3D classification, with a mask encompassing the cytoplasmic domains of KdpB. After non-uniform refinement and reconstruction, the resulting density maps were examined and compared with previous structures in the protein data bank, thus associating each class with a particular conformational state. These maps were then used as references for a series of hetero-refinement jobs to maximize the homogeneity of each particle set. After the sorting and recovery scheme outlined in Sfig. 1, non-uniform refinement was used to generate cryoSPARC density maps at resolutions shown in that figure. The E1∼P·ADP density map is included in the figure, even though it was described in our previous publication (9).

Although these maps displayed excellent resolution for the membrane domains, the cytoplasmic domains of KdpB were relatively disordered, reflecting innate flexibility and consequent structural heterogeneity. To improve the resolution of these cytoplasmic domains, we developed a workflow in RELION v5 (34). After importing the individual particle sets, 2D classification was used to remove a small fraction of bad particles. An initial reference was derived either from the cryoSPARC map (after noise normalization) or from a 3D initial reference job in RELION. An initial 3D auto-refine job did not include a mask, but after completion, was then continued using a mask derived from a fitted atomic model of the entire KdpFABC complex. Continuation of the auto-refine job retains previously determined particle orientations and proceeds with relatively fine sampling intervals and ranges. Once finished, the auto-refine job was continued again, this time with a mask corresponding to the KdpB subunit. Finally, the job was continued a third time with a mask corresponding only to the cytoplasmic A-, N- and P-domains from KdpB (Sfig. 2). Because different masks were used in each case, the reported resolutions are not directly comparable. Nevertheless, the cytoplasmic domains, and specifically the catalytic site involving Asp307 in the P-domain, the nucleotide binding site in the N-domain, and the conserved TGES^162^ sequence in the A-domain, were considerably better resolved. The maps shown in Fig. 1 represent a hybrid of the maps from cryoSPARC and RELION.

Initial atomic models were built by fitting published structures into the individual density maps (7LC3 for E1 conformations and 7BGY for E2 conformations) and then rebuilding each using COOT. For well resolved regions such as the transmembrane domains, the sharpened map from cryoSPARC was used. For other regions, sharpened and unsharpened maps from both cryoSPARC and RELION were used. For particular poorly ordered regions such as A- and N-domains from the E1apo structure, the respective domains were fitted as a rigid body. These starting models were then refined with PHENIX (35) using the sharpened cryoSPARC map for less mobile regions (typically transmembrane and P-domain) and the unsharpened RELION map for the cytoplasmic regions. Initially, these refinements were performed with truncated atomic models, whereas at later stages the full model was used with coordinate restraints applied to the alternate regions with a sigma of either 0.5 or 1.0. For analysis of the tunnels, we used the CavitOmiX plugin (v. 1.0, 2022, Innophore GmbH) to the PyMol Molecular Graphics System (Schrödinger, LLC) with a probe size of 0.8 and a grid spacing of 0.4. This plugin uses a modified LIGSITE algorithm to delineate cavities within proteins (36). Most figures were created with CHIMERAX (37), though surface coloring according to the local resolution was done with CHIMERA (38).

### Activity assays

Methods for measuring ATPase and transport activity have been described in recent publications (9, 33). Briefly, ATPase activity was measured using a coupled enzyme assay (39) in a standard buffer containing 75 mM TES-Tris pH 7.3, 7.5 mM MgCl_2_, 2.5 mM ATP and 150 mM KCl. To assess K^+^ stimulation as shown in Fig. 4, the buffer composition was 25 mM TES-Tris, 2.5 mM MgCl_2_, and 1 mM ATP in the absence of added KCl; both ATP and MgCl_2_ were reduced to minimize K^+^ contamination bound in those reagents (9).

To assess the effect of lipids on ATPase activity of detergent-solubilized preparations, lipid stock solutions were prepared by first using Argon gas to evaporate chloroform and produce a thin film, which was further dried under vacuum for 1-2 h, resuspended and dissolved in aqueous solutions containing DM to final lipid concentration of 8.3 mg/mL. For DOPA, DOPC, and DOPE 15% DM was used, whereas other lipids were dissolved in 5% DM. Lipids were then added to the ATPase assay solutions in 20 µL increments followed by ∼1 min measurement of activity. All lipids were obtained from Avanti Polar Lipids (Alabaster, AL).

For measuring ATPase activity of reconstituted samples, we first added minimal amounts of DM to open the vesicles, thus exposing internally facing proteins to ATP and preventing a buildup of K^+^ inside the vesicles. The concentration of DM was determined empirically for each proteolipsome preparation by titration of DM during measurement of ATPase (Sfig. 7); the highest activity from each titration curve was reported. To compare activities from different preparations, we quantified the reconstitution efficiency by measuring the density of the KdpA band in Coomassie-stained SDS-PAGE gels; detergent-solubilized preparation of WT KdpFABC was used as a standard.

Transport was measured by solid-supported membrane electrophysiology as previously described (9, 33). Briefly, KdpFABC was reconstituted into liposomes with a 4:1 weight ratio of POPC and DOPA with a final lipid concentration of 5 mg/mL and a lipid to protein weight ratio of 5:1. After passivating sensors according to the manufacturer’s recommendation, the SURFE^2^RN1 (Nanion Technologies, Livingston, NJ) was used to record currents. Buffers contained 50 mM HEPES-Tris, pH 7.1, 5 mM MgSO_4_, 50 mM K_2_SO_4_ with 300 µM ATP added for activation of transport. As in previous work (9), transport activity was quantified by recording current at 1.25 s, thus avoiding transient binding signals that are sometimes seen with mutant samples. As with ATPase activity, transport currents from each preparation were normalized according to the reconstitution efficiency determined by SDS-PAGE analysis in Sfig. 7.

### MST experiments

KdpFABC mutants were labeled with Alexa Fluor 488 dye (Invitrogen Life Technologies) by adding 2.0 µL of dye from a 16 mM stock solution in DMSO to 100 µL of protein at a concentration of 1–2 mg/mL in 100 mM NaCl, 25 mM Tris pH 7.5, 10% glycerol, 1 mM TCEP, and 0.15% DM (SEC buffer). Excess free dye was removed by size-exclusion chromatography using SEC buffer. Lipids were prepared by drying 5 mg of lipids under Ar gas and dissolving them in SEC buffer containing 5% DM at a concentration of 8 mM. For titrations, lipid concentrations were varied by serial dilution and then mixed with 100 nM protein solution at a 1:1 ratio by volume. The titration mixtures were incubated for 30 minutes and loaded into standard treated capillaries for measurement on a Monolith NT.115 instrument (NanoTemper Technologies, South San Francisco, CA). Data were acquired using LED power settings of 20–40% and medium MST power. Three to four independent titrations were collected and analyzed with MO.Affinity Analysis software v2.3 using an MST on-time of 20 s. Dissociation constants (Kd) were obtained by fitting the data to a binding curve based on the law of mass action:

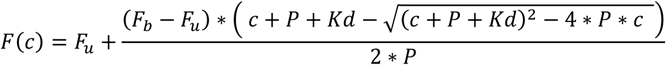

Where c is concentration of lipid species, P is the concentration of Kdp protein and F(c) is the normalized fluorescence signal measured at a given concentration. In addition to Kd, unknowns for this equation are normalized fluorescence for protein in the bound state (F_b_) and normalized fluorescence for protein in the unbound state (F_u_). Typically, this data is replotted, substituting normalized fluorescence with fraction bound (Sfig. 9).

### Mass Spectrometry

Samples were suspended in methanol/chloroform (2:1) and incubated at 37°C for 30 minutes to denature proteins. Chloroform and water were added, the samples were vortexed, and phase separation was achieved by centrifugation. The lower phase was collected, dried under nitrogen, and re-dissolved in 0.1 ml chloroform/methanol (1:1). Lipids were analyzed by LC-ESI-MS/MS on a QExactive HF-X instrument coupled directly to a Vanquish UHPLC (Thermo Fisher Scientific, Waltham, MA, USA). An aliquot of 7 µl was injected into a Restek Ultra C18 reversed-phase column (Restek Corporation, Bellefonte, PA, USA; 100×2.1 mm; particle size 3 µm) that was kept at a temperature of 50°C. Chromatography was performed with solvents A and B at a flow rate of 0.15 mL/min. Solvent A contained 600 ml acetonitrile, 399 ml water, 1 ml formic acid, and 0.631 g ammonium formate. Solvent B contained 900 ml 2-propanol, 99 ml acetonitrile, 1 ml formic acid, and 0.631 g ammonium formate. The chromatographic run time was 40 minutes, changing the proportion of solvent B in a non-linear gradient from 30 to 35% (0-2 min), from 35 to 67% (2-5 min), from 67 to 83% (5-8 min), from 83 to 91% (8-11 min), from 91 to 95% (11-14 min), from 95 to 97% (14-17 min), from 97 to 98% (17-20 min), from 98 to 100% (20-25 min), and from 100 to 30% (25-26 min). For the remainder of the run, solvent B remained at 30% (26-40 min). The mass spectrometer was operated in negative ion mode. The spray voltage was set to 4 kV and the capillary temperature was set to 350°C. MS1 scans were acquired in profile mode at a resolution of 120,000, an AGC target of 1e6, a maximal injection time of 65 ms, and a scan range of 200-2000 m/z. MS2 scans were acquired in profile mode at a resolution of 30,000, an AGC target of 3e6, a maximal injection time of 75 ms, a loop count of 7, and an isolation window of 1.7 m/z. The normalized collision energy was set to 30 and the dynamic exclusion time to 31 s. For lipid identification and quantitation, data were analyzed by the software LipidSearch 5.1.8 (Thermo Fisher Scientific, Waltham, MA, USA). The general database was searched with a precursor tolerance of 3 ppm, a product tolerance of 10 ppm, and a product intensity threshold of 1.0%.

## Acknowledgements

Screening of cryo-EM samples was performed at NYU Langone Health’s Cryo-Electron Microscopy Laboratory (RRID:SCR_019202), which is partially supported by the Laura and Isaac Perlmutter Cancer Center Support Grant NIH/ NCI P30CA016087 and NIH Grant R01NS108151. Cryo-EM data collection was supported by NIH grant R24GM154185 and performed at the Pacific Northwest Center for Cryo-EM (PNCC) with assistance from Sean Mulligan. The authors acknowledge support from Novo Nordisk Foundation grant NNF24OC0088380, from the European Research Council (ERC) under the European Union’s Horizon 2020 research and innovation program (grant agreement no. 101000936) and from the Danish National Research Foundation under the grant DNRF190 entitled Center for active transport of plant hormones (Plant-PATH) to BPP. The project was also supported by the National Institutes of Health grant R35GM144109 to DLS and grant R01HL166418 to MS.

## Notes

### Competing Interest Statement

The authors have declared no competing interest.

