## Supplemental Figures for "Lipids are essential for potassium transport by KdpFABC from E. coli"

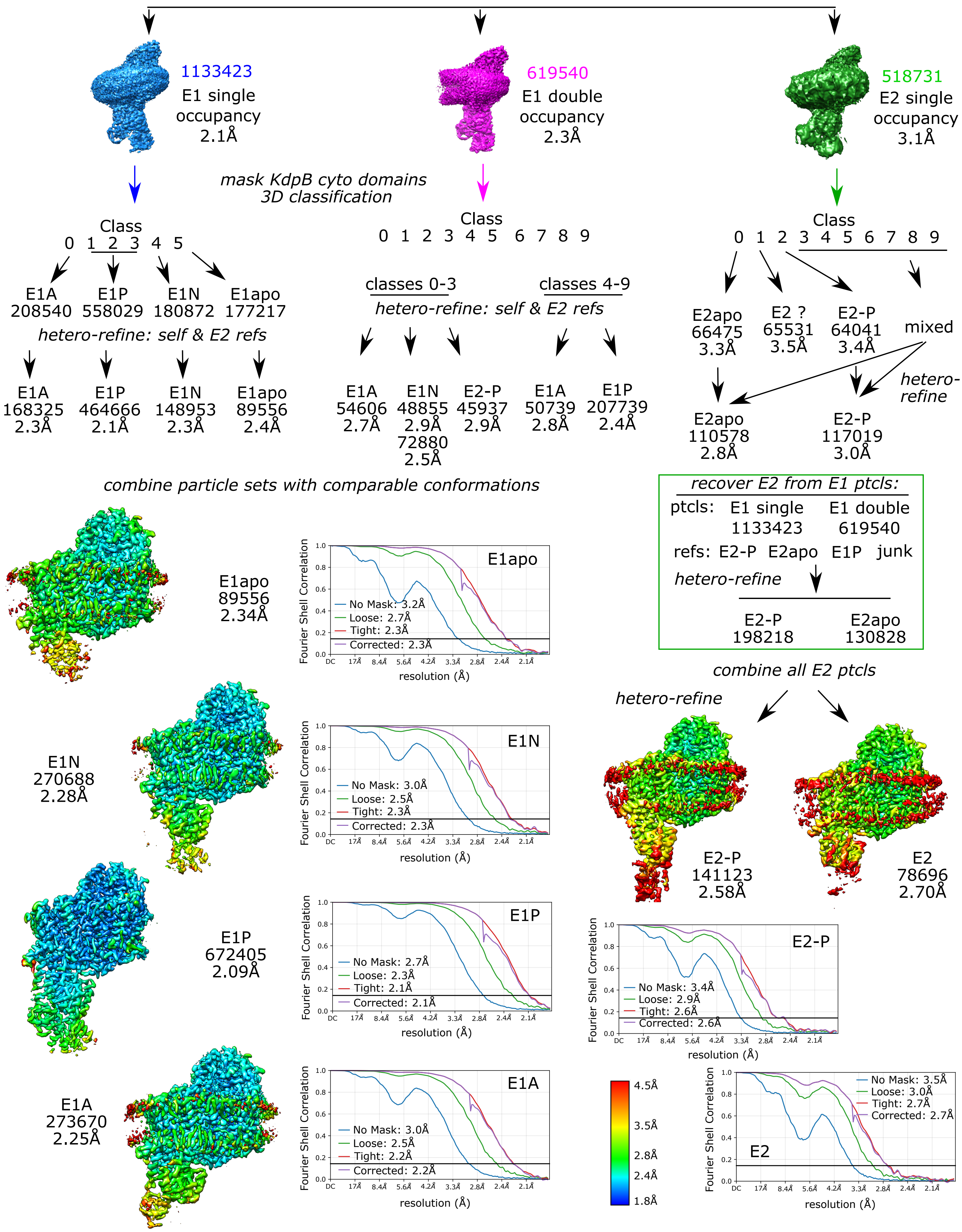
1133423
E1 single occupancy
2.1Å
619540
E1 double occupancy
2.3Å
518731
E2 single occupancy
3.1Å
mask KdpB cyto domains
3D classification
Class
0 1 2 3 4 5
E1A 208540
E1P 558029
E1N 180872
E1apo 177217
hetero-refine: self & E2 refs
E1A 168325 2.3Å
E1P 464666 2.1Å
E1N 148953 2.3Å
E1apo 89556 2.4Å
Class
0 1 2 3 4 5 6 7 8 9
classes 0-3
classes 4-9
hetero-refine: self & E2 refs
E1A 54606 2.7Å
E1N 48855 2.9Å 72880 2.5Å
E2-P 45937 2.9Å
E1A 50739 2.8Å
E1P 207739 2.4Å
Class
0 1 2 3 4 5 6 7 8 9
E2apo 66475 3.3Å
E2 ? 65531 3.5Å
E2-P 64041 3.4Å
mixed
hetero-refine
E2apo 110578 2.8Å
E2-P 117019 3.0Å
combine particle sets with comparable conformations
recover E2 from E1 ptcls:
ptcls: E1 single 1133423
E1 double 619540
refs: E2-P E2apo E1P junk
hetero-refine
E2-P 198218
E2apo 130828
combine all E2 ptcls
hetero-refine
E1apo 89556 2.34Å
E1apo
No Mask: 3.2Å
Loose: 2.7Å
Tight: 2.3Å
Corrected: 2.3Å
Fourier Shell Correlation
resolution (Å)
E1N 270688 2.28Å
E1N
No Mask: 3.0Å
Loose: 2.5Å
Tight: 2.3Å
Corrected: 2.3Å
E1P 672405 2.09Å
E1P
No Mask: 2.7Å
Loose: 2.3Å
Tight: 2.1Å
Corrected: 2.1Å
E1A 273670 2.25Å
E1A
No Mask: 3.0Å
Loose: 2.5Å
Tight: 2.2Å
Corrected: 2.2Å
E2-P 141123 2.58Å
E2 78696 2.70Å
E2-P
No Mask: 3.4Å
Loose: 2.9Å
Tight: 2.6Å
Corrected: 2.6Å
E2
No Mask: 3.5Å
Loose: 3.0Å
Tight: 2.7Å
Corrected: 2.7Å
4.5Å
3.5Å
2.8Å
2.4Å
1.8Å

### Supplemental Figure 1. Cryo-EM processing pipeline using cryoSPARC

Outline of the processing steps in cryoSPARC used to derive six structures from a single sample. Our previous report (9) described initial processing of ~50,000 images, which resulted in 3 main classes, shown at the top of this figure (E1 single occupancy, E1 double occupancy and E2 single occupancy). The occupancy refers to the number of KdpFABC complexes occupying the nanodisc; although two copies are visible in class averages from double occupancy particles, the second copy becomes averaged out during refinement. Masked 3D classification of these three classes, based on A-, N- and P-domains of KdpB, was crucial in segregating the conformational states. Once suitable references for each state were obtained, heterogeneous refinement jobs were employed to clean up particle sets and to recover mis-assigned particles from earlier classification jobs. The final density maps are shown for each state, colored according to local resolution (key shown at the bottom). The plots show Fourier Shell Correlation for each refinement with final resolution determined from the corrected, masked maps according to the gold standard threshold of 0.143. The density map for E1P has been previously reported (9) and deposited in the protein data bank with accession numbers 9OC4 and EMD-70308.

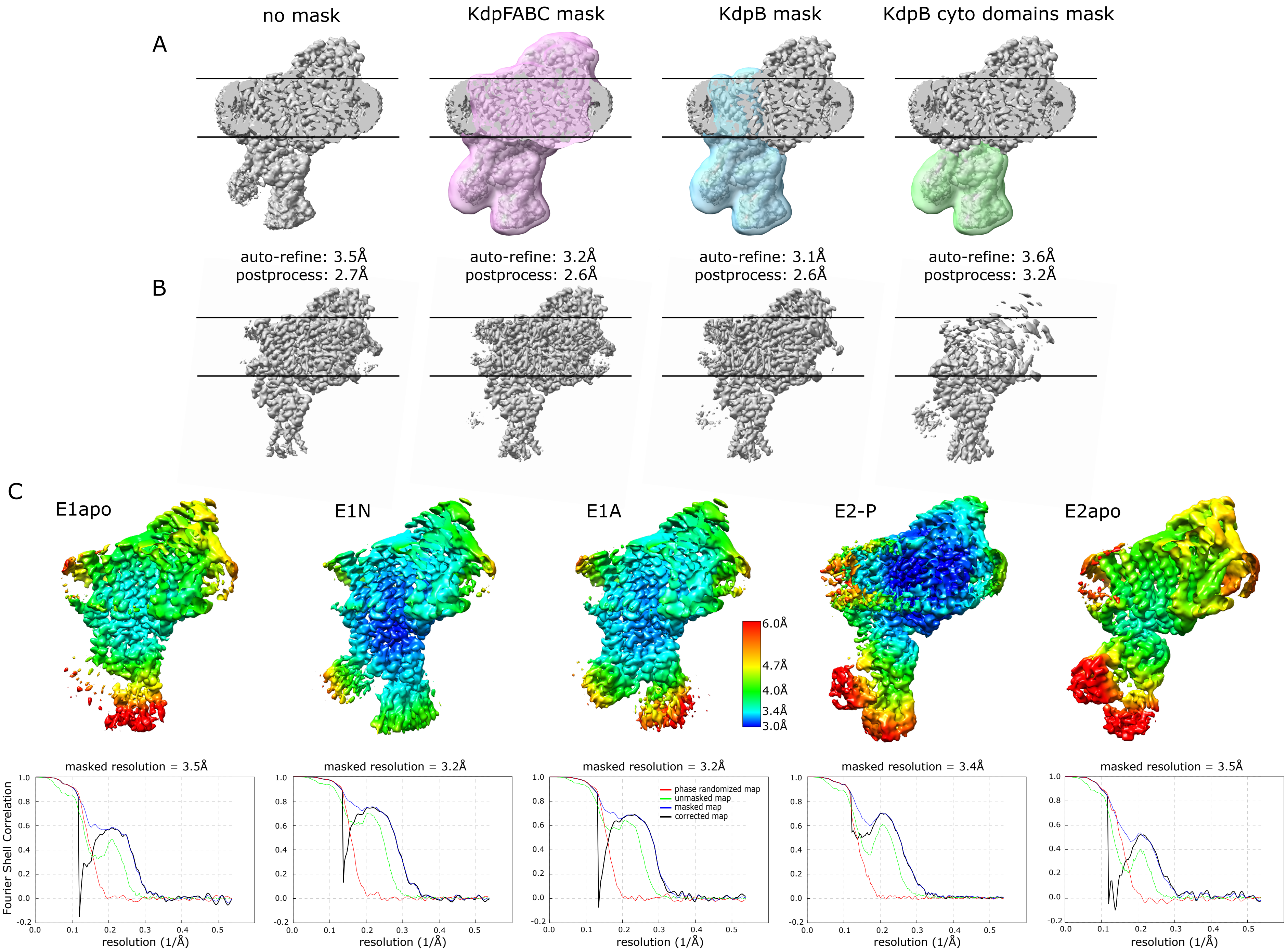
no mask
KdpFABC mask
KdpB mask
KdpB cyto domains mask
A
auto-refine: 3.5Å
postprocess: 2.7Å
auto-refine: 3.2Å
postprocess: 2.6Å
auto-refine: 3.1Å
postprocess: 2.6Å
auto-refine: 3.6Å
postprocess: 3.2Å
B
C
E1apo
E1N
E1A
E2-P
E2apo
6.0Å
4.7Å
4.0Å
3.4Å
3.0Å
masked resolution = 3.5Å
masked resolution = 3.2Å
masked resolution = 3.2Å
masked resolution = 3.4Å
masked resolution = 3.5Å
Fourier Shell Correlation
resolution (1/Å)
phase randomized map
unmasked map
masked map
corrected map

### Supplemental Figure 2. Cryo-EM processing pipeline using RELION

To better resolve the flexible cytoplasmic domains of KdpB, a masked refinement strategy was employed using RELION. To start, final particle sets from Sfig. 1 were imported into RELION. (A) A consecutive series of refinements were run starting with no mask and then gradually focusing the refinement on the cytoplasmic domains of KdpB (green mask). (B) The resulting maps from the E1N dataset at step are shown together with their respective resolutions from post processing jobs. Note that focus on the cytoplasmic domains caused the transmembrane domains to become blurred. (C) Final maps from focused refinement colored according to local resolution (key shown in the middle). Fourier Shell Correlation plots are shown along the bottom with resolutions derived from the corrected masked map at a threshold of 0.143.

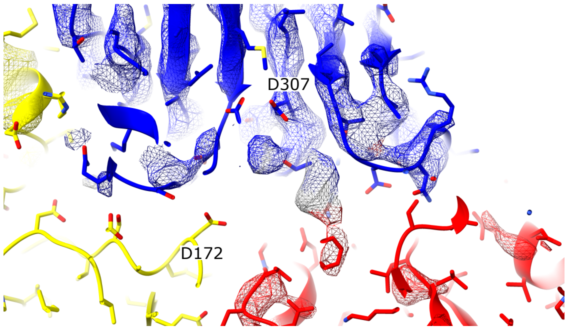


E1

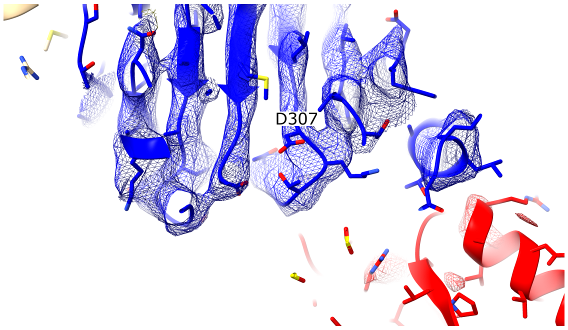


E2

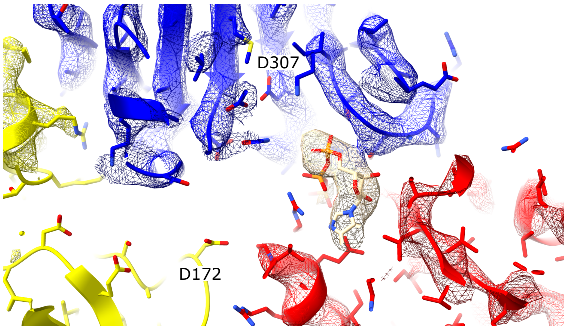


E1·ATP

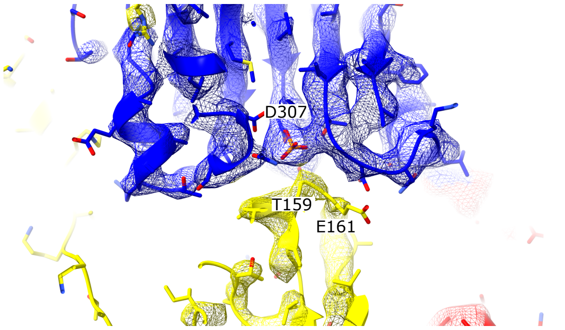


E2-P

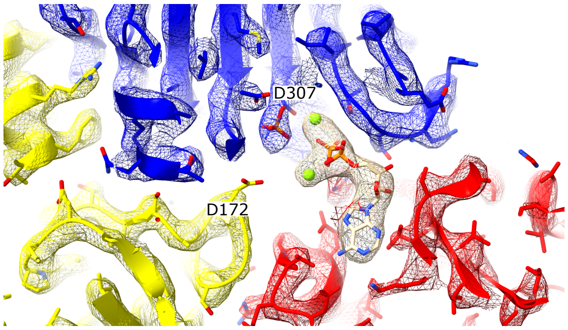


E1-P·ADP

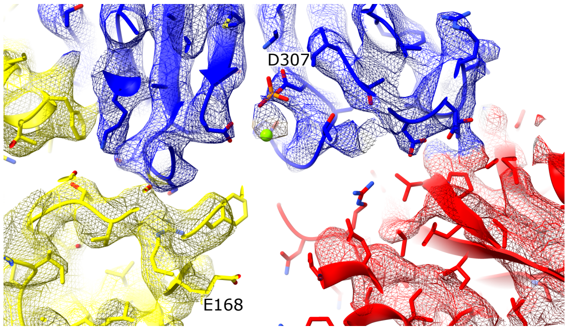


E1-P

### Supplemental Figure 3. Map densities at the catalytic site of KdpB

Gallery of images showing the catalytic site in all states. In the E1~P·ADP state, ADP is bound by the N-domain (red) along with two $Mg^{2+}$ ions (green) and Asp307 in the P-domain carries a phosphate. In the E1·ATP state (referred to as E1N in Sfigs. 1 and 2), a nucleotide is bound to the N-domain but Asp307 is not phosphorylated. In E1~P (referred to as E1A in Sfigs. 1 and 2), Asp307 is phosphorylated, but nucleotide is not present in the N-domain. In E2-P, Asp307 is phosphorylated and the conserved $TGES^{162}$ loop in the A-domain (yellow) is nearby, ready to hydrolyze the aspartyl phosphate. In E2, Asp307 is not phosphorlated, and both N- and the A-domains have moved away from the site. In E1 (referred to as E1apo in Sfigs. 1 and 2), the cytoplasmic domains are positioned similarly to E1~P·ADP, but Asp307 is not phosphorylated and the N-domain is empty.

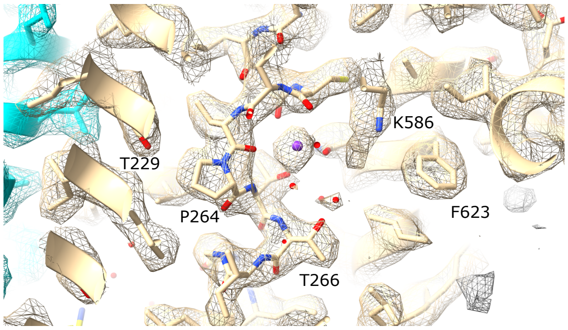


E1

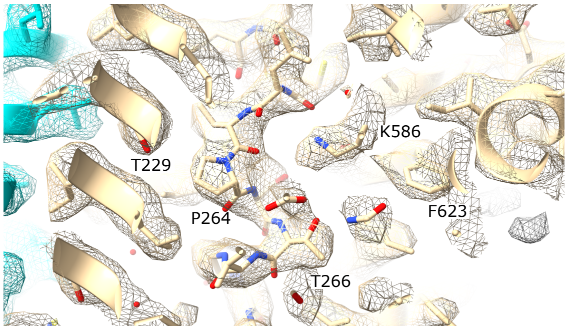


E2

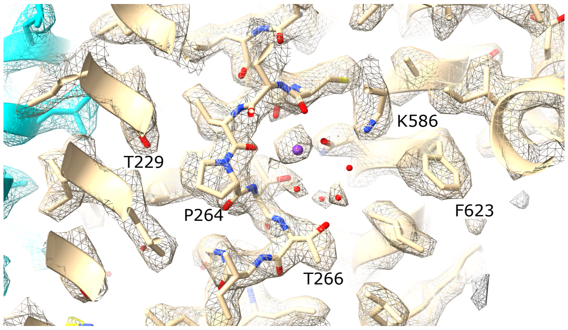


E1·ATP

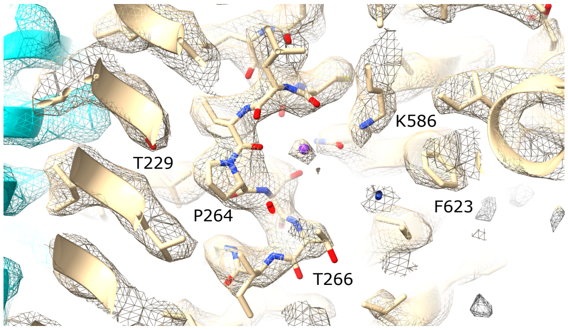


E2-P

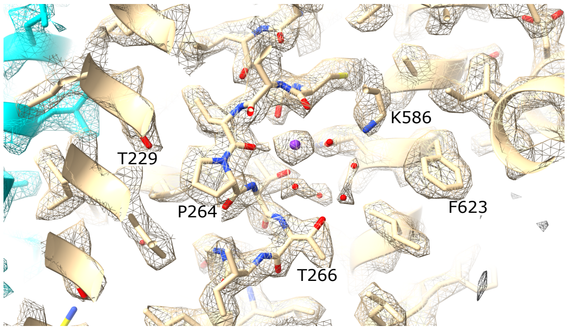


E1-P·ADP

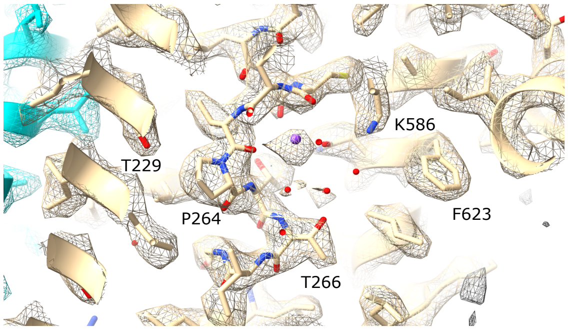


E1-P

### Supplemental Figure 4. Map densities at the CBS

Gallery of images showing the CBS in all states. This site is essentially unchanged in all E1 states, with a strong density at the $K^+$ binding site and Lys586 retracted. In E2-P, this conformation remains unchanged; in E2, however, movements of M5 cause Lys586 to swing into the site and thus displace $K^+$.

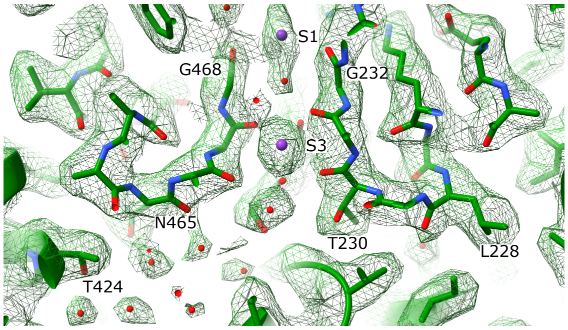


E1

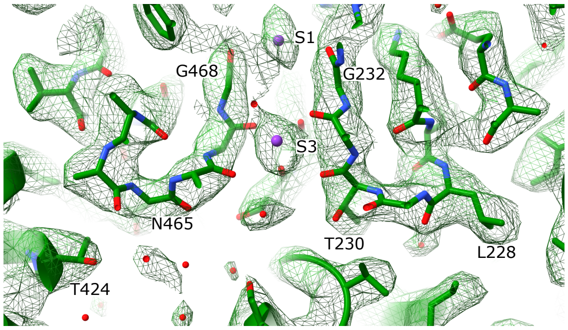


E2

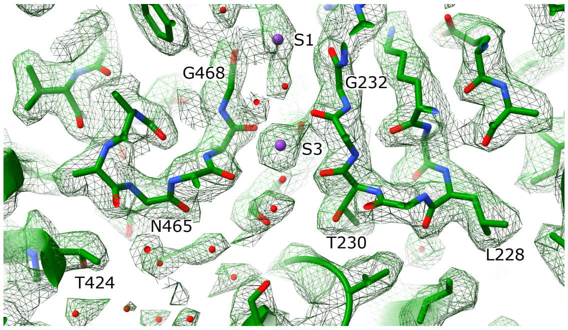


E1·ATP

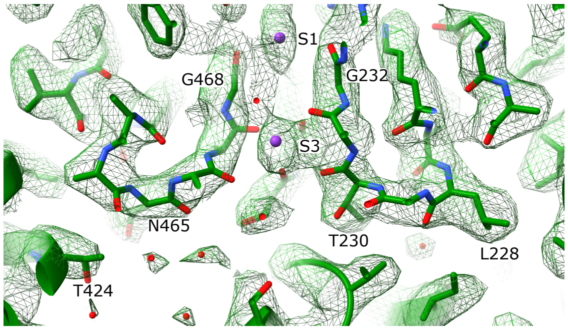


E2-P

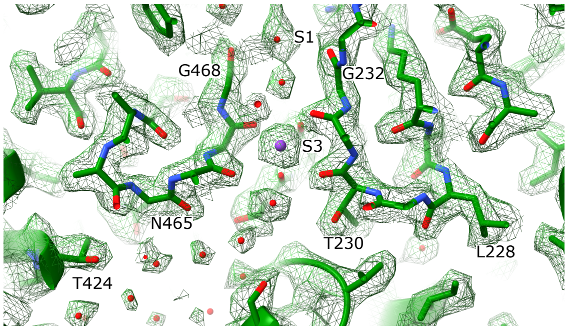


E1-P·ADP

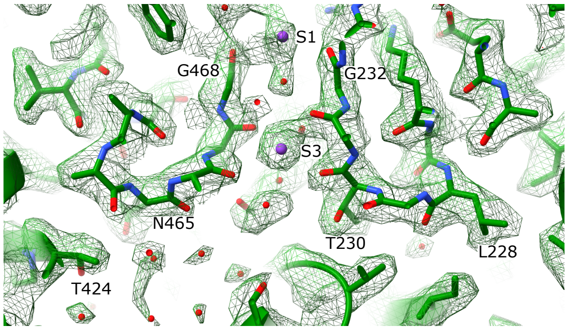


E1-P

### Supplemental Figure 5. Map densities at the selectivity filter

Gallery of images showing that the selectivity filter remains static throughout the reaction cycle, as reflected by the low RMSD values for KdpA and KdpC in Table 2. Very strong density is observed at the S3 site of the SF in all structures, indicating the presence of $K^+$ at this primary binding site. Most of the structures also have strong density at the S1 site at the entrance to the SF, which was therefore also modeled as $K^+$. Weaker densities at S2 and S4 and throughout the intersubunit tunnel were modeled as water.

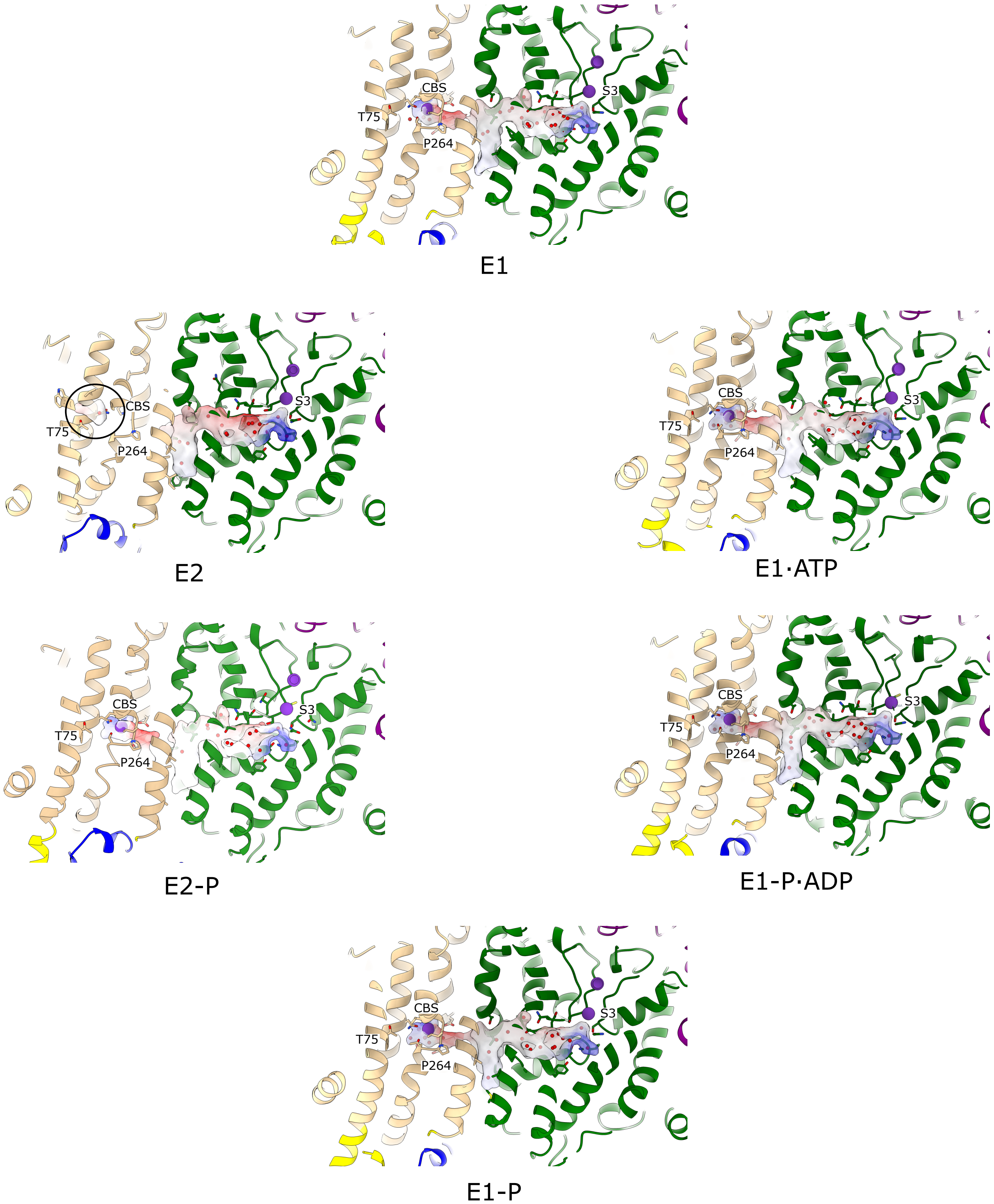
CBS
T75
P264
S3
E1
CBS
T75
P264
S3
E2
CBS
T75
P264
S3
E1·ATP
CBS
T75
P264
S3
E2-P
CBS
T75
P264
S3
E1-P·ADP
CBS
T75
P264
S3
E1-P

### Supplemental Figure 6. Dynamics of the intramembrane tunnel

Gallery of images showing the intramembrane tunnel running between S3 sites of the selectivity filter in KdpA (green) and the CBS in KdpB (tan) next to Pro265. The tunnel has been colored according to electrostatic potential of the surrounding protein: color scheme as in Fig. 3e. The tunnel is pinched at the interface between KdpA and KdpB in E2-P. In E2, the cavity at the CBS collapses completely and a new cavity appears next to Thr75 (black circle), which was previously shown to be the $K^+$ release site (9).

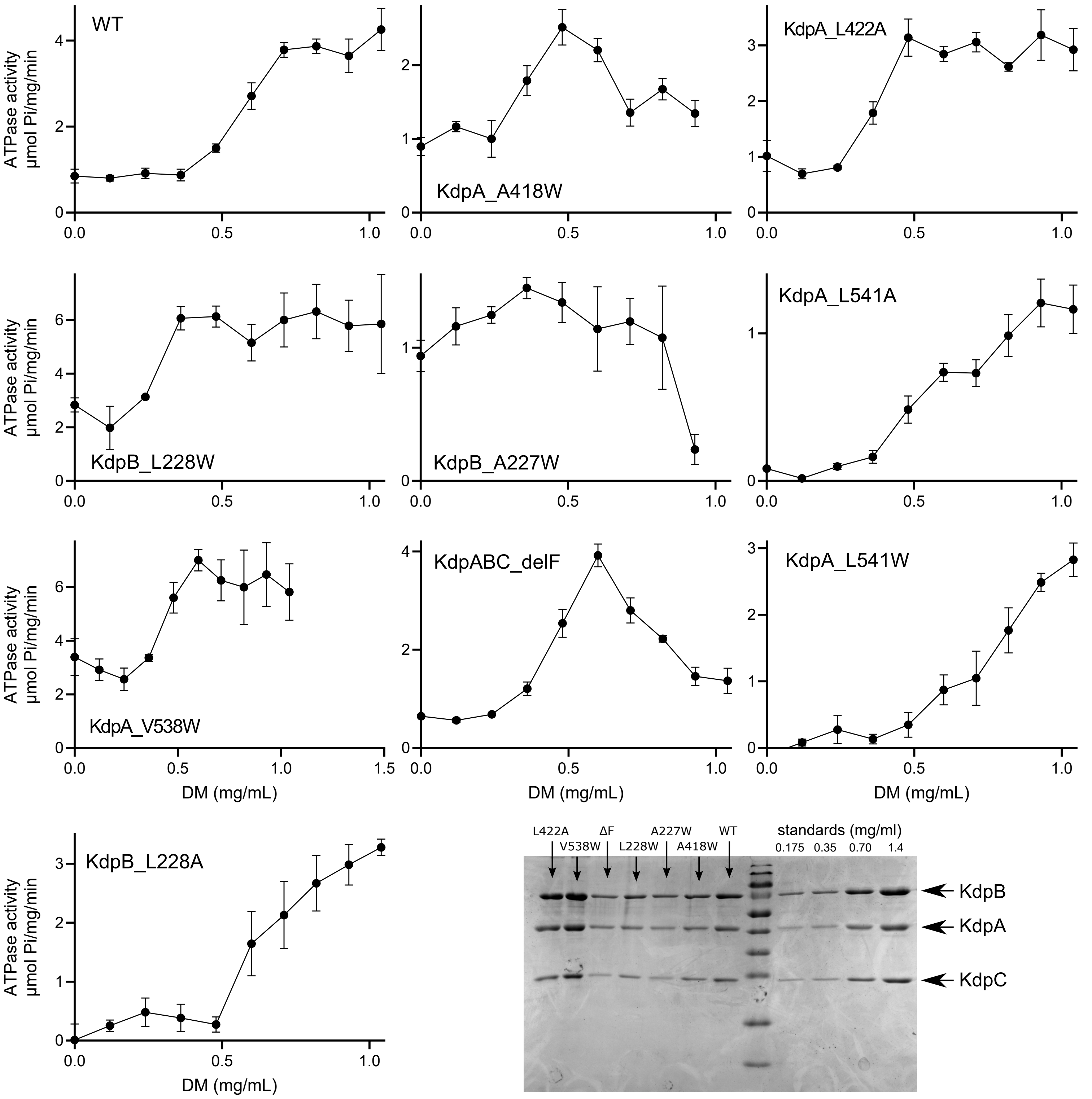
WT
KdpA_A418W
KdpA_L422A
KdpB_L228W
KdpB_A227W
KdpA_L541A
KdpA_V538W
KdpABC_delF
KdpA_L541W
KdpB_L228A
ATPase activity
μmol Pi/mg/min
DM (mg/mL)
L422A
V538W
ΔF
L228W
A227W
A418W
WT
standards (mg/ml)
0.175
0.35
0.70
1.4
KdpB
KdpA
KdpC

### Supplemental Figure 7. Detergent treatment of proteoliposomes

To measure ATPase activities from reconstituted proteoliposomes, DM was added to allow ATP to reach inward facing molecules and to prevent buildup of membrane potential and internal $K^+$, which would otherwise inhibit activity. For each titration, the maximal ATPase activity was plotted in Fig. 4b. Note that some mutants were inhibited by excess detergent, indicating their sensitivity to delipidation. SDS-PAGE (lower right) was used to quantify the amount of protein in each reconstituted preparation, which was then used to normalize the associated ATPase and transport activity in Fig. 4. Error bars correspond to SEM based on three technical replicates.

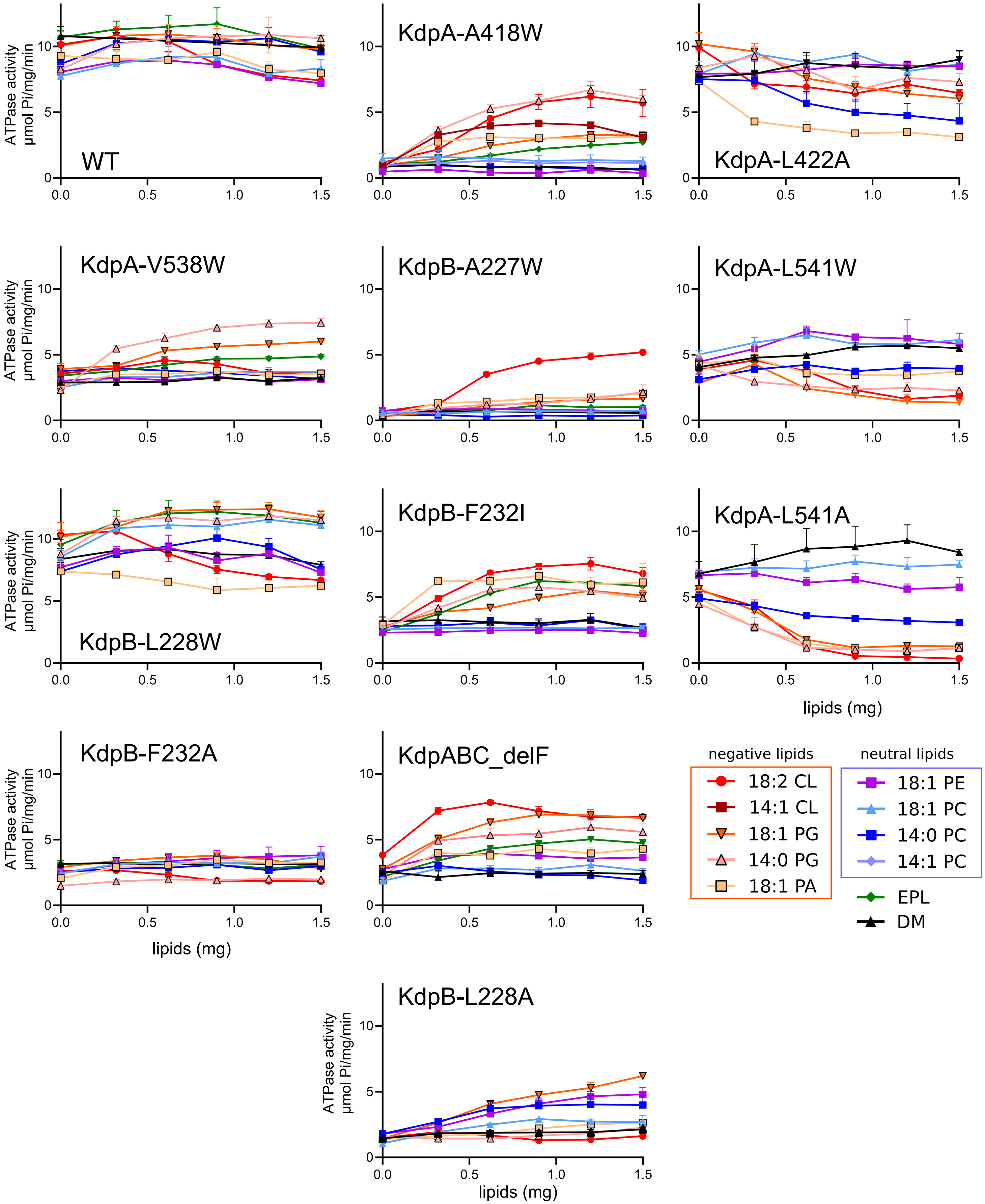
ATPase activity µmol Pi/mg/min
WT
KdpA-A418W
KdpA-L422A
KdpA-V538W
KdpB-A227W
KdpA-L541W
KdpB-L228W
KdpB-F232I
KdpA-L541A
KdpB-F232A
KdpABC_delF
KdpB-L228A
lipids (mg)
negative lipids
18:2 CL
14:1 CL
18:1 PG
14:0 PG
18:1 PA
neutral lipids
18:1 PE
18:1 PC
14:0 PC
14:1 PC
EPL
DM

### Supplemental Figure 8. Effect of lipid on ATPase activity of detergent solubilized samples

ATPase activities of detergent solubilized samples are plotted as a function of added lipid for each mutant, where lines simply connect the individual data points. To derive values for Fig. 5, data from DOPG, DOPE and DOPC were fitted either by linear regression or by the Michaelis-Menton equation in order to calculate an averaged value of activity at 1.5 mg. Error bars represent SEM for three technical replicates.

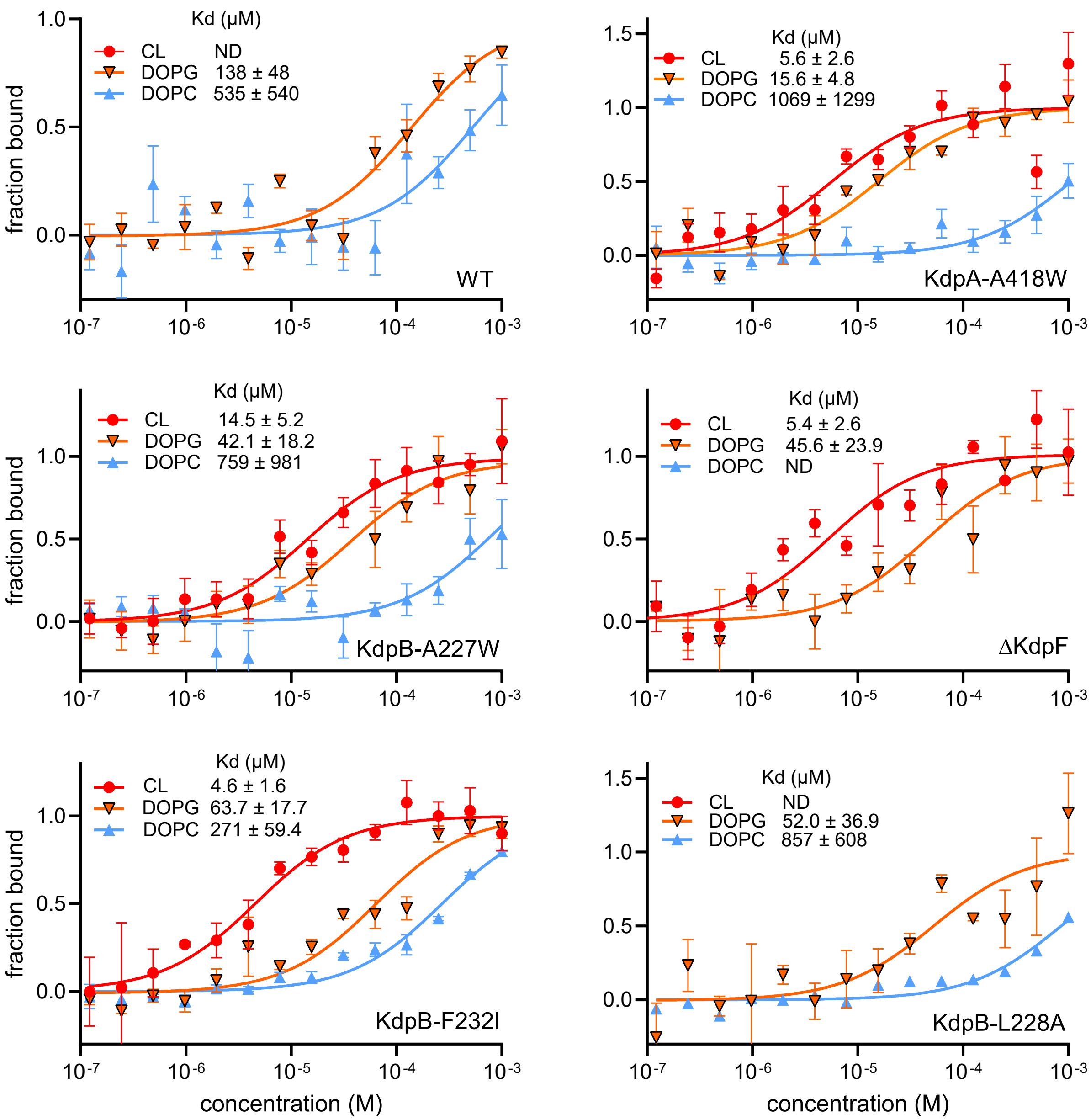
Kd (µM)
CL ND
DOPG 138 ± 48
DOPC 535 ± 540
WT
fraction bound
Kd (µM)
CL 5.6 ± 2.6
DOPG 15.6 ± 4.8
DOPC 1069 ± 1299
KdpA-A418W
Kd (µM)
CL 14.5 ± 5.2
DOPG 42.1 ± 18.2
DOPC 759 ± 981
KdpB-A227W
Kd (µM)
CL 5.4 ± 2.6
DOPG 45.6 ± 23.9
DOPC ND
ΔKdpF
Kd (µM)
CL 4.6 ± 1.6
DOPG 63.7 ± 17.7
DOPC 271 ± 59.4
KdpB-F232I
Kd (µM)
CL ND
DOPG 52.0 ± 36.9
DOPC 857 ± 608
KdpB-L228A
concentration (M)

### Supplemental Figure 9. MST measurements of lipid binding

Titration curves for MST measurements of lipid binding to detergent solubilized preparations. These data were derived by the manufacturers software and were then fitted with Michaelis-Menton equation to calculate the binding constant (note $K_d$ ± SEM is reported here whereas $K_a$ is reported in Fig. 5). In some cases, the software failed to detect binding and these cases are indicated by "ND" in the legend. Error bars represent SEM from three technical replicates.

### Supplemental Movie 1

Morph between E1~P state and E2-P state showing structural changes depicted in Fig. 6.

### Supplemental Movie 2

Morph between E2-P state and E2 state showing structural changes depicted in Fig. 6.
